# Optimal sampling design for spatial capture-recapture

**DOI:** 10.1101/2020.04.16.045740

**Authors:** Gates Dupont, J. Andrew Royle, Muhammad Ali Nawaz, Chris Sutherland

## Abstract

Spatial capture-recapture (SCR) has emerged as the industry standard for estimating population density by leveraging information from spatial locations of repeat encounters of individuals. The precision of density estimates depends fundamentally on the number and spatial configuration of traps. Despite this knowledge, existing sampling design recommendations are heuristic and their performance remains untested for most practical applications. To address this issue, we propose a genetic algorithm that minimizes any sensible, criteria-based objective function to produce near-optimal sampling designs. To motivate the idea of optimality, we compare the performance of designs optimized using three model-based criteria related to the probability of capture. We use simulation to show that these designs out-perform those based on existing recommendations in terms of bias, precision, and accuracy in the estimation of population size. Our approach allows conservation practitioners and researchers to generate customized and improved sampling designs for wildlife monitoring.

## Introduction

The need for conservation managers and practitioners to obtain reliable estimates of population size (Williams et al., 2002) has driven the rapid development of data collection and estimation methods. Capture-recapture (CR), and more recently, spatial capture-recapture (SCR; Efford, 2004; Borchers and Efford, 2008) methods were developed specifically for this purpose and are now routinely applied in ecological research. Concurrently, SCR methods estimate detection, space use, and density by analyzing individual encounter histories while explicitly incorporating auxiliary information from the spatial organization of encounters (Efford, 2004; Royle et al., 2014). Despite widespread adoption and rapid method development, recommendations about spatial sampling design have received relatively little attention and are arguably heuristic.

The effects of sampling design have been investigated for both CR (Dillon and Kelly 2007; Bondrup-Nielsen 1983) and SCR methods (discussed below). While CR methods aim to balance the number of captures and the number of recaptures, SCR requires a third consideration, the spatial pattern of individual encounter histories. The ability to reliably estimate density is directly related to these considerations: the number of captured individuals *n* is the sample size; the number of recaptures is directly related to the baseline detection probability, *g*_0_; and the number and spatial distribution of recaptures are directly related to the spatial scale parameter, *σ*. Therefore, improving sampling design has great potential to increase the quality of the data and the precision of parameter estimates.

Several simulation studies evaluating SCR designs have shown that inference is robust to the spatial configuration of traps, as long as some minimum requirements are met: the trap spacing must not be too large relative to individual space use in order to reliably estimate *σ*, but the array must not be too small such that too few individuals are exposed to capture (Sollmann et al., 2012; Sun et al., 2014; Wilton et al., 2014; Efford and Boulanger, 2019; Tobler and Powell, 2013). Repeated illustrations of this trade-off have lead to recommendations that trap spacing should be approximately two times *σ*, which maximizes accuracy and minimizes bias of abundance estimates (Sollmann et al., 2012; Efford and Fewster, 2013; Efford and Boulanger, 2019). While most of this research has focused on uniform grids, simulation has also shown that clustered designs can outperform uniform designs (Efford and Fewster, 2013; Sun et al., 2014), particularly for heterogeneously distributed populations (Efford and Fewster, 2013; Wilton et al., 2014). In summary, the idea of optimal sampling design for SCR remains poorly understood beyond these few, basic recommendations. In particular, it is unclear whether existing design heuristics generally hold for spatially-varying density patterns, or in highly-structured landscapes where recommended regular trapping arrays can not be accommodated, and guidance of generating clustered designs is lacking.

Generally speaking, sampling design for SCR can be conceived as a problem of selecting a subset of all possible trap locations that maximizes some SCR-relevant objective function. Here we develop an analytical framework that directly addresses this challenge. Our approach generates a near-optimal sampling design with respect to some appropriately defined objective function and information about available resources (traps), a set of all possible trap locations, and information about SCR model parameters. To motivate the idea of optimality, we use simulation to compare the performance of existing recommendation to designs optimized using three model-based criteria related to current thinking about the relationship between data quality and estimator bias and precision. We explore design performances for scenarios where we vary the spatial coverage of traps, the landscape geometry, and deviations from uniform spatial distribution of individuals.

## Methods

### The standard SCR model

Typically, SCR models have two model components: a spatial model of abundance describing the distribution of individuals characterized by the center of their home range (hereby referred to as an activity center), and a spatial model of detection that relates encounter rates to the distance between the activity center and a trap (e.g., a camera trap). The most basic form assumes a uniform prior for the distribution of activity centers, *s*_*i*_:

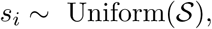

where *𝒮*, referred to as the state-space, describes all possible locations of activity centers. To facilitate analysis, *𝒮* is represented as a uniform grid of points representing the centroids of equal-sized pixels. All individuals within the region, *N*, are exposed to capture resulting in the observation of *n* individuals and hence *n*_0_ = *N* − *n* unobserved individuals.

While several formulations of the encounter model exist, we use, without loss of generality, a half-normal encounter model that describes encounter probability as a decreasing function of distance from an individual’s activity center *s*_*i*_:

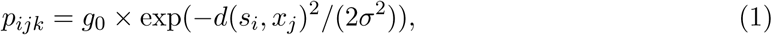

where *p*_*ijk*_ is the probability of detection of individual *i* with activity center *s*_*i*_ at trap *j* during sampling occasion *k*; *d*(*s*_*i*_, *x*_*j*_) is the distance between the activity center *s*_*i*_ and the trap *x*_*j*_, and *g*_0_ and *σ* are the baseline encounter probability and spatial scale parameters, respectively.

### Model-based objective functions

From Equation 1, we can use values of *g*_0_ and *σ* (e.g., from the literature or estimates from a pilot study), to compute the probability that an individual with an activity center *s*_*i*_ is detected in *any* trap in an array *𝒳*, which we denote as 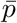:

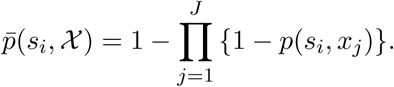

The corresponding marginal probability of not being encountered is thus: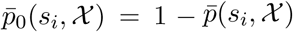. Taking the average over all *G* activity center locations in the landscape *𝒮*, we can compute the marginal probability of encounter:

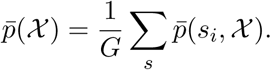

We can also compute the probability of being captured in exactly one trap:

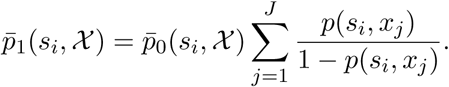

Finally, the marginal probability of being encountered at more than one trap, i.e., of a spatial recapture is:

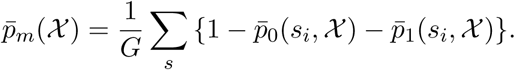

Given that the precision of SCR density estimates depends on the total number of individuals captured, *n*, and the number of spatial recaptures, *m* (Efford and Boulanger, 2019; Royle et al., 2014) 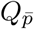 and 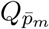 represent logical criteria for optimizing SCR designs (Royle et al. 2014, Chapter 10). Herein lies one of our novel contributions: we suggest three design criteria: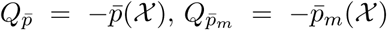, and 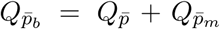. Importantly, if approximate values of the SCR parameters, *g*_0_ and *σ*, are available, these objective functions can be evaluated analytically for any number and configuration of traps, providing a metric for efficient identification of optimal SCR designs.

### Optimization method

We applied a genetic algorithm (GA) to the task of finding a design that minimizes any criterion, noting that optimality here is with respect to the defined criteria, and in the context of the GA is ‘near-optimal’ (see Appendix S1 & Goldberg, 1989). The GA is a random search algorithm which produces multiple generations of solutions, where subsequent generations retain characteristics of top performing solutions from the previous generation. Generations are produced until converging on a near-optimal solution is achieved. Wolters (2015) adapted the algorithm to solve a *k-of-n* problem which describes concisely the challenge of the SCR sampling design: the selection of some number of traps, *k*, in a landscape of *n* possible locations according to some objective function. We provide a detailed description of the general GA, the *k-of-n* adaptation, and our implementation in the R package `oSCR` in Appendix S1 and Appendix S4.

Conceptually, minimizing the space-filling objective function 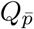 maximizes the expected sample size *n*. In contrast, minimizing 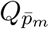 prioritizes the exposure of individuals to more than one trap and should maximize the number of spatial recaptures *m*. The third criteria, 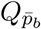, attempts to balance 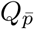 and 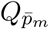.

### Design constraints

We were primarily interested in evaluating the performance of SCR designs produced by our framework under a range of biologically-realistic scenarios in an attempt to develop a more general understanding of how performance varies as a function of the following design constraints: *geometry*, defined as the shape of the study area and ease at which a regular square trapping grid can be deployed; *density pattern*, defined as the nature of departure from uniform distribution of individuals; and *effort*, defined as the number of traps available for the design.

#### Geometry

As has been typical in studies investigating SCR sampling designs, we begin using a square study area with complete accessibility and which lends itself to uniform trapping grids (the *regular area*, Figure 1). To replicate the design challenges posed when generating real-world designs, we also consider an *irregular area* (Figure 1). For this, we use one of the study areas that motivated this work: a large area in Northern Pakistan (3865 *km*^2^) that is the focus of a snow leopard *(Panthera uncia)* camera trapping study, but that has several logistical challenges that determine accessibility (i.e., remoteness, private property, altitude, and slope). To define the complete region of the state-space, we used a 3*σ* buffer around the trapping extent. The regular area is represented by 24 × 24 landscape with a resolution of 0.5 units, the irregular study area is represented by 89.85 × 133.04 landscape with a resolution of 1.73 units, for a total of 2304 cells in each of the geometries (Figure 1). While these two state-spaces differ in absolute terms, we insured comparability in relative terms by the definition of area-specific sigma (see below).

**Figure 1.**
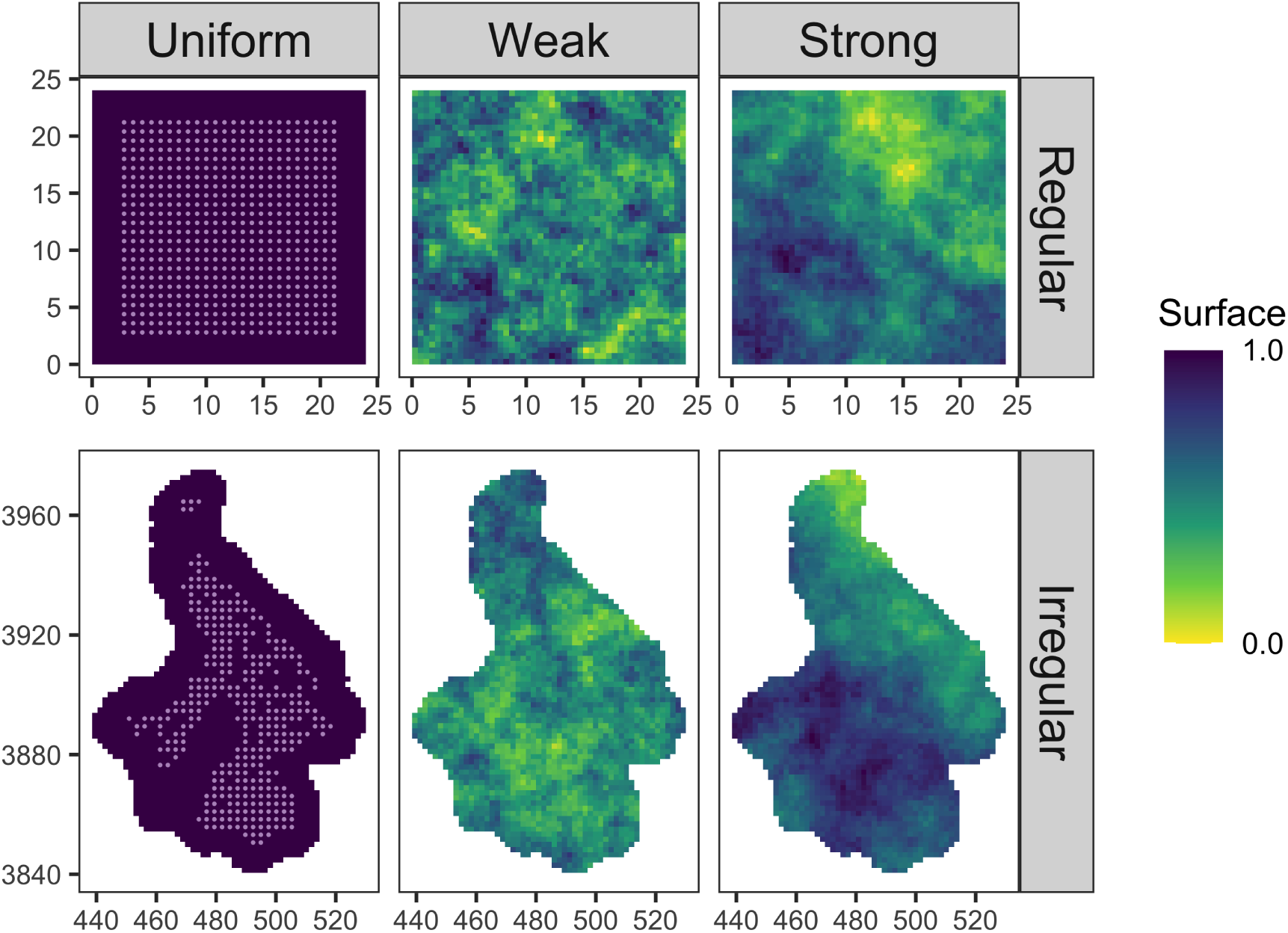
Simulation structure. Here we show all possible trap locations overlaid on the uniform landscape for the regular (top) and irregular (bottom) study area geometries alongside a single realization of two (weak: middle, strong: right) of the three (uniform not shown) landscape covariates. For the regular geometry, we tested 12 designs each. For the irregular geometry, we tested 9 designs each. This makes for a total of 63 scenarios.

#### Density pattern

Existing investigations of SCR sampling designs typically assume a homogeneous distribution of individuals (but see Efford and Fewster, 2013). Here we formally test the adequacy of designs under specific violations of this assumption. We consider three spatial density patterns: a uniform and two spatially-varying. To generate non-uniform density patterns, we simulated landscapes defined by a parametric Gaussian random field that allows for specification of the degree and range of spatial autocorrelation. Gaussian random fields were generated using the R package, `NLMR` (Sciaini et al., 2018). The values of the simulated landscape were scaled from 0 to 1 and individual activity centers distributed according to the following cell probabilities:

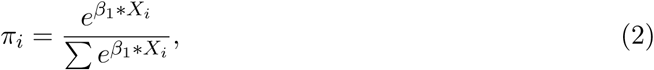

where *X*_*i*_ is the scaled landscape value at pixel *i* and *β*_1_ is defined as 1.2 to represent a weak but apparent density pattern. The two inhomogeneous density patterns differ in the scale of spatial autocorrelation. For consistency, we defined this distance in relative terms to the length of the longest side of the state-space: 6% for a *weak* density pattern that produces a patchy landscape, and 100% for a *strong* density pattern produces a landscape with a more contiguous gradient (see Figure 1 for a single realization of the density patterns). Using these three density patterns allows us to evaluate designs through a full range of biological realism, with uniform and strong density patterns representing the polar ends of reality, and the patchy landscape representing the most realistic sampling scenario.

### Design generation

Designs were generated using fixed values of *g*_0_ and *σ* (see below), a set of potential trap locations, and the number of traps that are available to deploy. It is assumed that the user has knowledge or access to data on information approximate values of SCR parameters, would be able to produce a set of all potential sampling points, and would have some idea of resources (traps) available. For the regular area, we generated 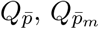, and 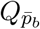 designs for each of the three levels of effort where there was no restriction on where traps could be placed. In addition, we generated a regular 2*σ* design for comparison. For the irregular area in the mountains of Pakistan, we generated only criteria-based designs at each of the three levels of effort (Figure 2). In this case, areas known to be too remote, too high altitude, or too steep to be accessed were removed from the set of potential trap locations. Mirroring real design challenges faced by managers, it was not practical to generate a 2*σ* grid for the irregular area, and therefore it is not included. This full scenario analysis resulted in a total of 21 designs; 12 designs for the regular area (the three optimized and the 2*σ* design), and 9 designs for the irregular area (optimized designs only).

**Figure 2.**
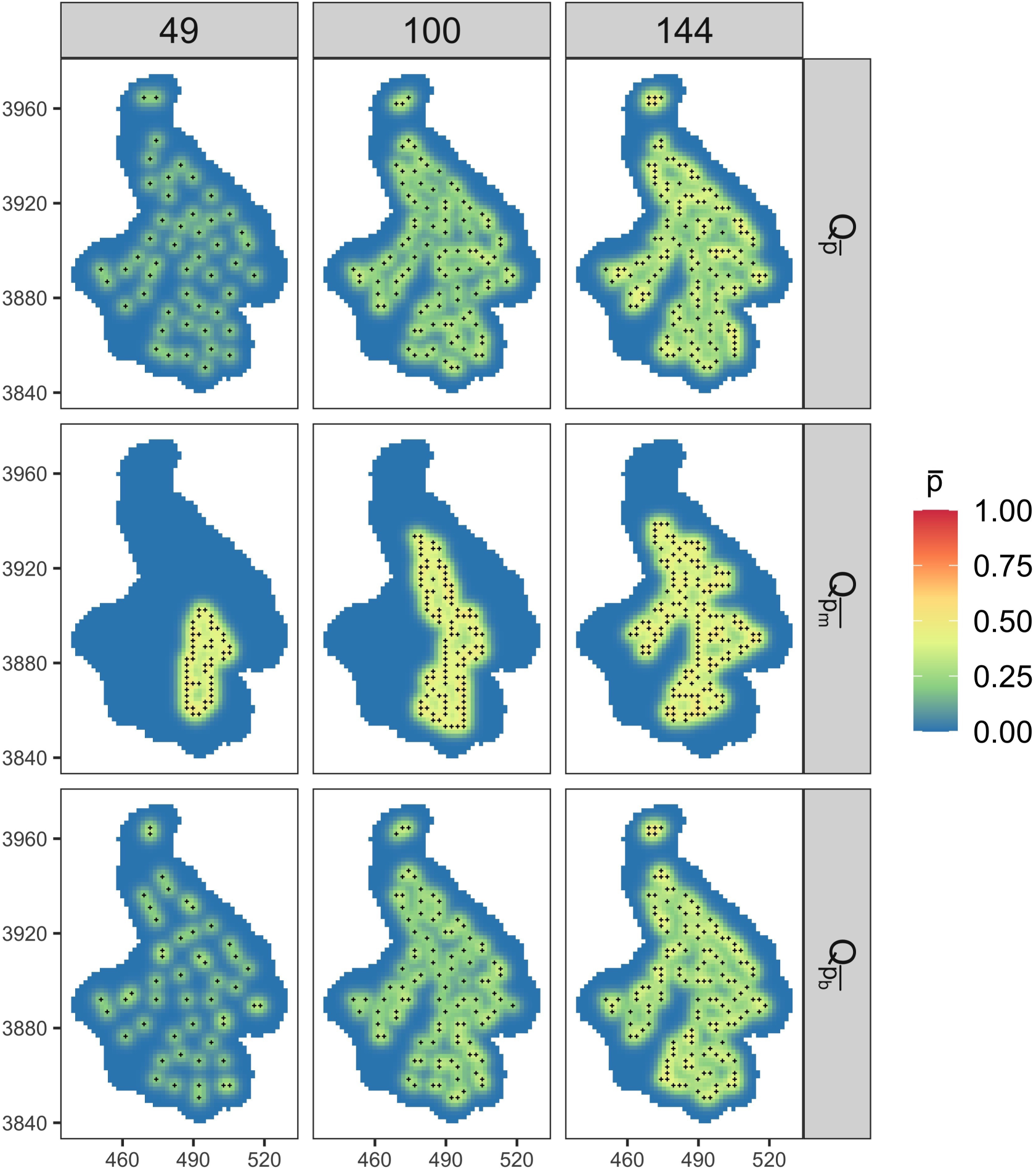
Irregular study area with designs generated using our new framework with three SCR-intuitive, model-based criteria (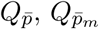, and 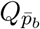), under three levels of effort. 144 traps represents the same number of traps as used to generate a full 2*σ* grid in a regular study area of the same area. 100 traps is nearly two-thirds as many traps, and 49 is nearly one-third as many traps. Each pixel of the state-space is colored according to the probability of capture, *p*, for an individual with an activity center at the centroid of the pixel.

### Evaluation by simulation

We exposed a population of *N* = 300 individuals to sampling via each of the 21 designs described above. We simulated encounter histories assuming proximity detectors and under the binomial encounter model encounter (Eq.1) with *g*_0_ = 0.2, *k* = 5. The two geometries differ in terms of their spatial units so area-specific *σ* values were chosen such that the number of home ranges required to fill the areas and achieve an equal density was equivalent: *σ*_*reg*_ = 0.80 and *σ*_*irreg*_ = 2.59. We simulated individuals according to the three density patterns described above (Eq.2), resulting in a total of 63 scenarios of interest (three density patterns for each of the 21 designs (Figure 2, Appendix S2).

For each scenario, we simulated 300 realizations of activity centers. Covariate surfaces were generated randomly using the same seed, again resulting in variation among simulations but consistency across scenarios. In some cases, the realization of activity centers did not provide at least one spatial recapture; we recorded the number of these *failure* and generated a new realization of activity centers until a single spatial recapture was obtained in order to proceed with model fitting. This only occurred for 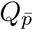 designs with minimum effort, and for less than 5% of the simulations.

We analyzed the resulting encounter history data using a null SCR model (*d*_***·***_) and, for spatially structured density scenarios, a density-varying model (*d*_*s*_). This allowed us to test if accounting for the landscape would improve bias and precision in parameter estimates. For each simulation, and each model, we retained estimates of *g*_0_, *σ*, and total abundance 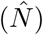.

We compared estimates of model parameters to the data-generating values in terms of bias (percent realtive bias, %RB), precision (coefficient of variation, CV), and accuracy (scaled root mean square error, SRMSE). All simulations were conducted in `R`, SCR models were fit using the package `oSCR` (Sutherland et al., 2019), and designs were generated using the `scrdesignGA()` function also in `oSCR` (detailed workflow provided in Appendix S3). Design generation and simulations were performed in R version 3.6.1 (R Core Team, 2019).

## Results

We first focus on relative bias. Encouragingly, under the regular-area, homogeneous-density scenario, designs generated using the genetic algorithm perform as well as existing 2*σ* recommendations, producing unbiased estimates of abundance for nearly all combinations of design and effort (Figure 3, Table 1). In the case of the irregular geometry with uniform density, 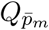 designs perform well for all levels of effort, but performance of 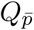 and 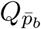 designs declines as the number of traps is reduced, a consequence of widely-spaced traps and consequently very few spatial recaptures (Figure 3, Table 1, Appendix S5, Appendix S6, Appendix S7).

**Table 1:** Percent relative bias of baseline detection (*g*_0_), space use (*σ*) and total abundance (EN) for each simulation scenario, varying: design criteria *(Design)*, landscape shape *(Geometry, Regular or Irregular)*, the number of traps *(Effort)*, and density patterns *(Density)*. We present results from null (*d*_*·*_) and varying density (*d*_*s*_) models.

| Effort | Density | Design | Regular |  |  |  |  |  | Irregular |  |  |  |  |  |
| --- | --- | --- | --- | --- | --- | --- | --- | --- | --- | --- | --- | --- | --- | --- |
| | | | $g_0$ | | $\sigma$ | | EN | | $g_0$ | | $\sigma$ | | EN | |
| | | | $d.$ | $d_s$ | $d.$ | $d_s$ | $d.$ | $d_s$ | $d.$ | $d_s$ | $d.$ | $d_s$ | $d.$ | $d_s$ |
| 49 | uniform | $2\sigma$ | 2.52 | — | -0.38 | — | 0.78 | — | — | — | — | — | — | — |
| | | $Q_{\bar{p}}$ | 0.82 | — | -1.00 | — | 7.27 | — | 2.27 | — | -1.84 | — | 8.34 | — |
| | | $Q_{\bar{p}_m}$ | 1.33 | — | -0.19 | — | 1.76 | — | 1.78 | — | -0.15 | — | 0.62 | — |
| | | $Q_{\bar{p}_b}$ | -0.61 | — | -2.06 | — | 13.32 | — | 2.53 | — | -4.11 | — | 17.90 | — |
| | weak | $2\sigma$ | 3.16 | 3.16 | -0.62 | -0.61 | -0.26 | -0.05 | — | — | — | — | — | — |
| | | $Q_{\bar{p}}$ | -0.58 | -0.58 | 0.20 | 0.25 | 5.70 | 5.75 | -1.51 | -1.51 | -1.11 | -1.07 | 9.93 | 9.89 |
| | | $Q_{\bar{p}_m}$ | 0.08 | 0.08 | 0.06 | 0.11 | 0.99 | 1.99 | 1.15 | 1.15 | -0.27 | -0.22 | 0.07 | 2.74 |
| | | $Q_{\bar{p}_b}$ | -2.73 | -2.73 | -2.36 | -2.16 | 16.12 | 14.83 | 0.19 | 0.19 | -1.48 | -1.46 | 13.68 | 14.09 |
| | strong | $2\sigma$ | 2.26 | 2.26 | -0.47 | -0.48 | 1.82 | 3.48 | — | — | — | — | — | — |
| | | $Q_{\bar{p}}$ | 1.84 | 1.84 | -0.75 | -0.78 | 6.43 | 6.55 | 1.18 | 1.18 | -0.27 | -0.32 | 5.80 | 6.17 |
| | | $Q_{\bar{p}_m}$ | 2.09 | 2.09 | -0.47 | -0.48 | 1.20 | 6.82 | 2.29 | 2.29 | -1.03 | -1.01 | 2.40 | 9.02 |
| | | $Q_{\bar{p}_b}$ | 0.99 | 0.99 | -3.47 | -3.41 | 14.54 | 14.10 | 2.75 | 2.75 | -3.32 | -3.26 | 15.13 | 15.14 |
| 100 | uniform | $2\sigma$ | 2.04 | — | -0.69 | — | 0.58 | — | — | — | — | — | — | — |
| | | $Q_{\bar{p}}$ | 2.42 | — | -0.61 | — | 0.90 | — | 1.42 | — | -0.77 | — | 2.11 | — |
| | | $Q_{\bar{p}_m}$ | -0.97 | — | 0.20 | — | 1.07 | — | 0.74 | — | -0.18 | — | 0.83 | — |
| | | $Q_{\bar{p}_b}$ | 0.07 | — | 0.05 | — | 1.12 | — | -0.15 | — | -0.51 | — | 2.55 | — |
| | weak | $2\sigma$ | -0.13 | -0.13 | 0.15 | 0.14 | -0.34 | -0.19 | — | — | — | — | — | — |
| | | $Q_{\bar{p}}$ | 0.61 | 0.61 | -0.27 | -0.29 | 0.95 | 0.98 | 0.97 | 0.97 | -0.48 | -0.49 | 1.82 | 1.89 |
| | | $Q_{\bar{p}_m}$ | 1.68 | 1.68 | -0.77 | -0.78 | -0.24 | 0.34 | -0.09 | -0.09 | 0.09 | 0.08 | 0.34 | 1.04 |
| | | $Q_{\bar{p}_b}$ | 1.07 | 1.07 | -0.16 | -0.18 | 0.01 | 0.03 | 1.23 | 1.23 | -0.30 | -0.27 | 1.06 | 1.11 |
| | strong | $2\sigma$ | 0.35 | 0.35 | -0.30 | -0.30 | 1.42 | 1.72 | — | — | — | — | — | — |
| | | $Q_{\bar{p}}$ | 0.18 | 0.18 | -0.93 | -0.95 | 2.89 | 3.12 | 1.07 | 1.07 | -0.46 | -0.49 | 0.93 | 1.40 |
| | | $Q_{\bar{p}_m}$ | 0.64 | 0.64 | -0.04 | -0.05 | 0.90 | 1.47 | 1.97 | 1.97 | -0.56 | -0.59 | -0.44 | 1.34 |
| | | $Q_{\bar{p}_b}$ | 0.60 | 0.60 | -0.43 | -0.43 | 1.36 | 1.44 | 0.21 | 0.21 | -0.05 | -0.06 | 0.40 | 0.80 |
| 144 | uniform | $2\sigma$ | 1.32 | — | -0.25 | — | 0.27 | — | — | — | — | — | — | — |
| | | $Q_{\bar{p}}$ | -1.06 | — | 0.28 | — | 1.53 | — | 0.72 | — | 0.08 | — | -0.27 | — |
| | | $Q_{\bar{p}_m}$ | 0.93 | — | -0.28 | — | 0.88 | — | 0.53 | — | 0.00 | — | 0.75 | — |
| | | $Q_{\bar{p}_b}$ | 0.35 | — | -0.07 | — | 0.90 | — | 2.12 | — | -0.77 | — | 0.72 | — |
| | weak | $2\sigma$ | 0.49 | 0.49 | -0.33 | -0.33 | 0.41 | 0.50 | — | — | — | — | — | — |
| | | $Q_{\bar{p}}$ | 0.64 | 0.64 | -0.24 | -0.25 | 0.44 | 0.47 | 0.61 | 0.61 | -0.20 | -0.20 | 0.50 | 0.51 |
| | | $Q_{\bar{p}_m}$ | 1.31 | 1.31 | -0.47 | -0.48 | -0.39 | -0.21 | 0.03 | 0.03 | 0.05 | 0.04 | 0.07 | 0.43 |
| | | $Q_{\bar{p}_b}$ | -0.02 | -0.02 | -0.32 | -0.33 | 1.00 | 0.98 | 0.77 | 0.77 | -0.25 | -0.26 | 0.93 | 0.92 |
| | strong | $2\sigma$ | 0.70 | 0.70 | -0.25 | -0.25 | 0.80 | 1.01 | — | — | — | — | — | — |
| | | $Q_{\bar{p}}$ | 1.35 | 1.35 | -0.31 | -0.32 | 0.32 | 0.47 | -0.13 | -0.13 | 0.21 | 0.19 | 0.33 | 0.66 |
| | | $Q_{\bar{p}_m}$ | 0.14 | 0.14 | 0.15 | 0.14 | 0.32 | 0.58 | 1.74 | 1.74 | -0.55 | -0.57 | -0.22 | 0.69 |
| | | $Q_{\bar{p}_b}$ | 1.18 | 1.18 | -0.19 | -0.20 | -0.03 | 0.14 | -0.59 | -0.59 | 0.12 | 0.09 | 0.20 | 0.62 |

**Figure 3.**
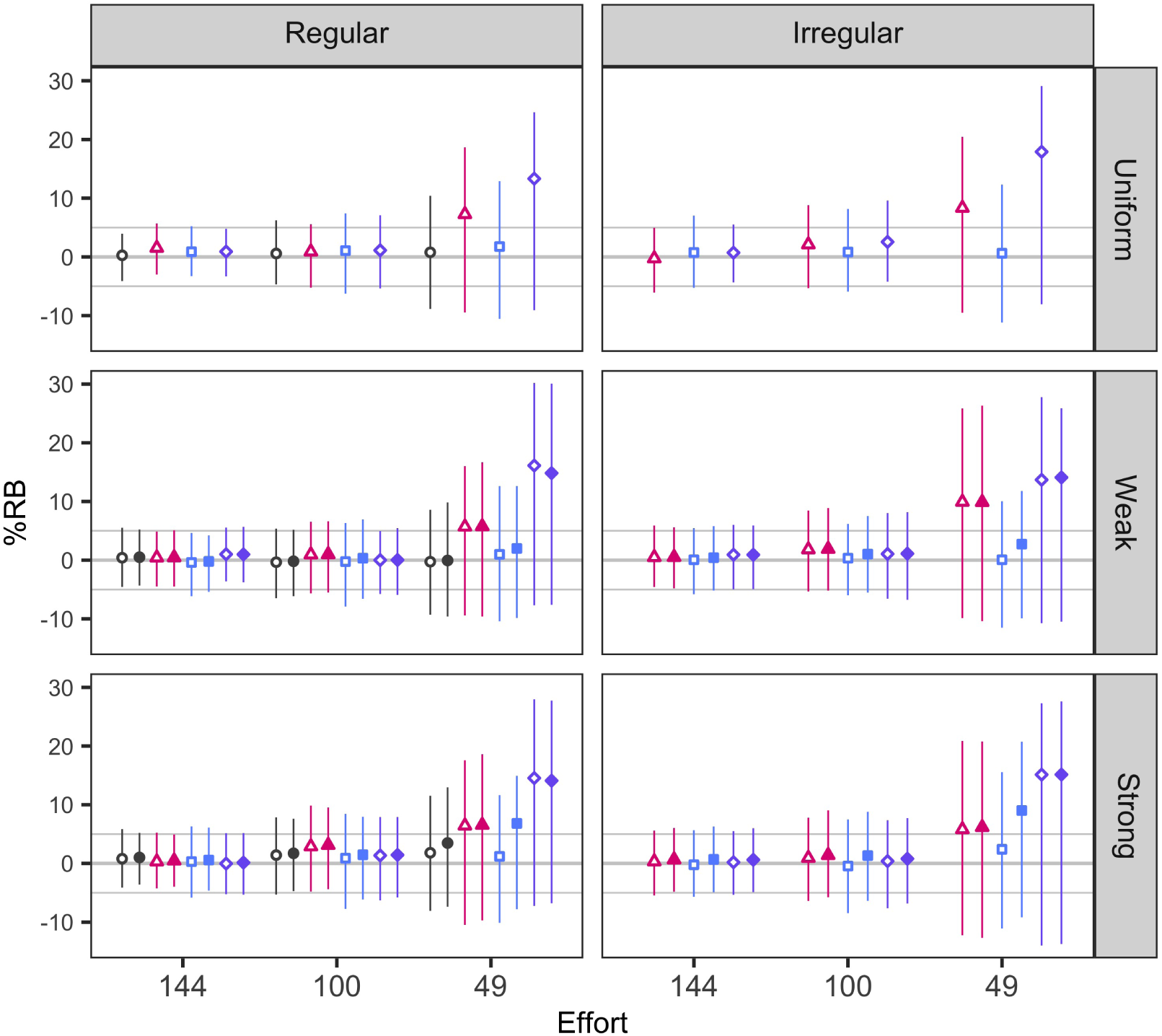
Percent relative bias (%RB) of estimates of total abundance from the three tested sampling designs under three levels of effort on three density surfaces within two geometries, where estimates are the result of one of two SCR models: density invariant (*d*_*·*_, open shapes) or density-varying (*d*_*s*_, closed shapes). The four designs – 2*σ*, 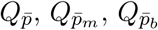– are represented by the four shapes: circles, triangles, squares, and diamonds, respectively. To illustrate estimator precision, vertical lines are 50% confidence intervals, noting that the 50% intervals are proportional to 95% intervals but offer a visual balance of bias and associated variance. The thick horizontal line represents no bias in estimates, with the thin horizontal lines representing an allowable amount of bias (*±* 5%).

For scenarios from the regular study area with inhomogeneous density, all designs produced unbiased estimates of abundance, generally. There is a slight bias (*±* 5%) introduced as the number of traps declines, even for the 2*σ* designs. However, this phenomenon is less apparent in 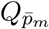 designs, suggesting improved performance. In the irregular study area, design performance is more dependent on the spatial structure of density. Once again, 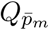 designs produced unbiased estimates, and 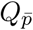 and 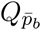 designs performed poorly with fewer traps (Figure 3, Table 1, Appendix S5, Appendix S6, Appendix S7).

Interestingly, explicitly including the landscape covariate governing spatial variation in density (i.e., *d*_*s*_ rather than *d*_***·***_) does not improve performance metrics for any of the designs in any scenario (Figure 3, Table 1), reinforcing the general opinion that SCR models are robust to misspecification of the density model. In fact, fitting the data-generating model for the inhomogeneous cases actually performs worse in low effort scenarios. This suggests that the low numbers of traps do not adequately represent the variation in the landscape, and therefore, the model is unable to reliably estimate the underlying landscape effect (Figure 3, Table 1).

Estimator precision and accuracy generally follow the same patterns as for the bias (Appendix S5 and Appendix S6, and Appendix S7, respectively). Design performance declines as effort decreases for all designs across every scenario. In the regular study area with uniform density, the 2*σ* and 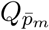 designs share similar levels of precision, while the 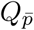 and 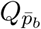 designs with minimal effort are less precise in comparison, with this pattern being magnified in the irregular area. Generally, there is a slight loss of precision in estimates across all designs, but this effect is less apparent for 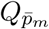 designs, which maintain their relative equivalency to the standard recommendation, including for the lowest level of effort (when considering comparison across geometries). In scenarios with inhomogenous density, both 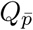 and 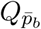 designs with minimum effort show precision that is obviously reduced using the null model. However, the density-varying model once again shows no noticeable improvement, and causes a decrease in precision for 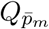 designs with the fewest traps.

Overall, designs generated using our proposed framework showed comparable performance to standard recommendations, and critically, these designs are robust to a variety of constraints that include effort, density signal, and geometry.

## Discussion

In this study, we develop a conceptual and analytical framework for generating near-optimal designs for SCR studies. We suggested three intuitive and statistically-grounded design criteria that can be optimized to produce candidate designs. We demonstrate that designs generated using our framework can perform at least as well as those based on existing heuristics, and further, that the generality and flexibility of our approach means it can be applied to any species or landscape according to logistics and available resources.

It is worth noting that the designs produced using this framework can be considered approximate in terms of specific location, and that the actual, finer-scale site-selection for traps can be informed by knowledge of the species’ biology and behavior (e.g., Fabiano et al., 2020). Further, while we develop this framework with camera traps in mind, this method can easily be applied to determine the general location of other non-invasive surveys, wherein the selection of a sampling location instead activates some other form of sampling effort (see Fuller et al. 2016; Sutherland et al. 2018). Importantly, the degree of sampling effort must be maintained among all selected sampling locations.

The designs we created using model-based criteria exhibit their own unique behaviors (Figure 2, Appendix S2). The 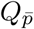 criteria generates space-filling designs to maximize the area covered and thereby the expected sample size of unique individuals. As more traps are added, the inner area becomes fully-saturated (such that it is insured that every possible home range will contain at least one trap), and the criteria instead focuses on selecting external traps that patrol the edge of the trapping extent in order to increase the probability of capture for individuals outside of that area. However, despite the benefit of increasing the sample size (*n* captured individuals), traps placed too distant from each other fail to generate important spatial recaptures. This is precisely the issue that propagated failures for both 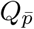 and 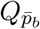 designs with minimum effort (Appendix S8).

In contrast, 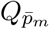 designs are space-restricting as a result of an inherent trade-off between increasing the number of individuals exposed to capture and having traps close together to insure captures at more than one trap. With fewer traps, however, the effective sampling area is markedly decreased (Figure 2), thereby reducing the sample size. This observation further motivated our evaluations of the designs for inhomogeneous density, which along with the reduced spatial coverage and hence non-representative sampling, is likely responsible for the bias observed in those scenarios, as well as the lower precision.

The 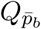 designs can best be described as “clustered space-filling” (Figure 2, Appendix S2), as this criteria aims to balance the objectives of 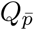 and 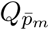, which it can do effectively when provided with a sufficient number of traps. However, as seen with 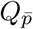 designs, the 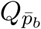 design performance suffers when too few traps are employed due to even larger distances between traps as a result of clustering, greatly reducing performance even beyond that of 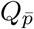.

More generally, these designs support previous recommendations while also providing new insights into sampling design for SCR. When full effort is possible in the regular area geometry, the 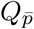 design fully saturates the trapping extent with some traps to spare in order to meet its objective, while 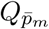 does not quite fill the trapping area (Figure 2, Appendix S2). Interestingly, the 2*σ* design falls somewhere between these two extents, likely striking an effective balance between the number of captures (as in 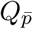) against the number of spatial recaptures (as in 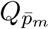), which we also see with 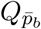 and similar to the effect described by Efford and Boulanger (2019). Despite these differences in spatial configuration, differences in design performance are mostly negligible (Figure 3, Table 1, Appendix S5, Appendix S6, Appendix S7).

As shown by Sun et al. (2014), incorporating trap clustering into sampling designs can be advantageous, as doing so allows for increased likelihood of spatial recaptures to facilitate estimation of the spatial scale parameter, *σ*. However, the clustered designs proposed by Sun et al. (2014) follow a regular pattern such that there are a limited number of levels of trap spacing, whereas the designs we generated result in a wider distribution of distances between traps. This shifts the importance away from a regular spatial structure of trap configuration to one that is decidedly irregular in order to gain better resolution of movement distances for estimating *σ*. This is especially useful knowledge and central to generating designs for irregular study areas. Interestingly, this results in designs with smaller effective sampling areas, suggesting that it might be better to reduce the total area covered by the design rather than focus on completely covering the area (within reason). A major insight here is that hierarchical clustering (the selection of approximately 2*σ*-spaced clusters of traps with further reduced within-cluster spacing) emerges naturally from the 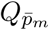 criterion, effectively formalizing the clustering heuristic proposed by Sun et al. (2014).

Our proposed criteria produced designs which perform well, yet there is scope for refinement. With a decrease in effective sampling area, the introduction of bias and imprecision in parameter estimates could be complicated further when the population being sampled has a stronger degree of spatial structuring than we tested here. Designs sampling only areas where individuals are concentrated will result in overestimates of population size and density relative to the whole study area, while those sampling away from concentrated areas will do just the opposite. This effect is particularly noticeable from the density-varying model (*d*_*s*_), which generally has relatively lower performance over the fully invariant model as it is including information from nearby traps sampling a landscape that is intrinsically spatially auto-correlated. Advancing this framework to generate designs that explicitly account for the spatial patterns in density as a function of a given landscape is clearly an area for further development, especially if the inferential objective is to estimate density-landscape relationships rather than density or total abundance.

Recently SCR sampling design for multi-species sampling has been considered, with some discussion on how the distribution of trap spacing can allow for better estimates for species with a variety of home range sizes (Rich et al., 2019). However, the design proposed for this purpose lacks a reproducible framework that can be generalized to any biological community. Alternatively, employing our framework for multi-species sampling could be a straightforward approach to this problem, with important implications for the use of SCR to be more easily applied for the study of ecological communities. Again, a highly appealing feature of our 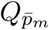 approach is the emergence of designs with much better distribution of trap spacing than under regular designs such as 2*σ* grids, ideal for sampling groups of species with varying spatial movement ecology.

We considered three criteria that are intuitive in the context of the performance trade off of sample size (*n*) and spatial recaptures (*m*). While intuitive, alternative criteria surely exist. For example, Efford and Boulanger 2019 propose an approximation of the variance of density which is related to *n* and *m*, and therefore can easily be formulated as an objective function to be optimized in the same way as 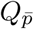 and 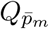. Indeed, the function `scrdesignGA()` is designed such that any user-defined objective functions can be used (e.g., Durbach *et al.* In review). We hope that this ability to simultaneously (and efficiently) generate and evaluate designs based on a variety of design criteria will motivate further research on SCR study design.

Our results show that designs obtained under our proposed criteria perform well relative to design heuristics and can be obtained efficiently as solutions to an optimization problem for arbitrary configurations of possible trapping locations and landscapes, unlike standard recommendations based on 2*σ* and cluster designs. Both CR and SCR studies are extremely expensive and require substantial effort to conduct, making it imperative that managers are provided with a method to select detector placement before deployment, such as the approach we have presented here. As a result, designs will produce a greater amount of expected information and will lead to more accurate estimates of parameters that describe biological populations of interest, which is critical to global conservation efforts, especially for low density and declining species that are of conservation concern but challenging to monitor.

## Supporting information

Metadata S1

## Acknowledgements

This work received support from Panthera, the Pakistan Snow Leopard and Ecosystem Protection Program, and the Snow Leopard Foundation. We thank the Sutherland Lab Group, especially Patricia Levasseur, as well as Katherine Zeller and Daniel Linden, for improving the manuscript. Any use of trade, product, or firm names is for descriptive purposes only and does not imply endorsement by the U.S. Government.

## Author contributions

CS, JAR, GD devised the study. CS and JAR wrote the functions for design generation. GD developed and conducted simulations. GD wrote the manuscript with contributions from all authors.

## Data availability

Metadata and code & data are available in Metadata S1 and Data S1.

