## Appendix S1 for "Optimal sampling design for spatial capture-recapture"

Appendix S1: Genetic algorithm details.

The design-generation function discussed here, `scrdesignGA()`, serves as a wrapper around the *k-of-n* genetic algorithm implemented by the R function `kofnGA()` (Wolters, 2015). The genetic algorithm that drives the design-generation process conducts a random search for solutions until it converges on a near-optimal solution. This algorithm is in the broader class of evolutionary algorithms, and following this, its components are named in similar terms.

The algorithm starts by generating a random set of possible solutions, which is the initial population of 'offspring' (in this context, designs). The size of the population is constant throughout the process, and is determined by the user via the *popsize* parameter. The offspring are then evaluated according to an objective function, resulting in a 'fitness' value for each offspring. Some proportion of the offspring are then allowed to 'breed' – determined via the *keepbest* parameter – which in effect is a mechanism that shares the 'genetic' material of the most fit offspring (i.e., the beneficial components) in order to make the next 'generation' of offspring. This process repeats for some number of generations, predefined by the *ngen* parameter. The *k-of-n* component of the specific genetic algorithm that we employ adds location-switching functionality that allows flexibility such that some number of traps, $k$, can be selected from $n$ possible trap locations, fitting neatly into the SCR design process.

While the three parameters mentioned above are important, we have found that only *ngen* is critical for parameter tuning of the algorithm. The algorithm should be parameterized to allow for a sufficient number of generations to reach (near-)convergence, which can be found via visual inspection by plotting the `scrdesignGA()` output object as illustrated in Appendix S4. Beyond that, and assuming the algorithm has converged on a (near-)optimal design, we have found that the genetic algorithm is not particularly sensitive to sensible values for the other two tuning parameters, *popsize* and *keepbest*, with the biggest differences relating to efficiency, which is case-specific and not critical overall.

See the main text of this manuscript for details regarding the required SCR components for design-generation, and for more details on the genetic algorithm we direct the reader to Wolters (2015), as cited in the main text.
