## Appendix S2 for "Optimal sampling design for spatial capture-recapture"

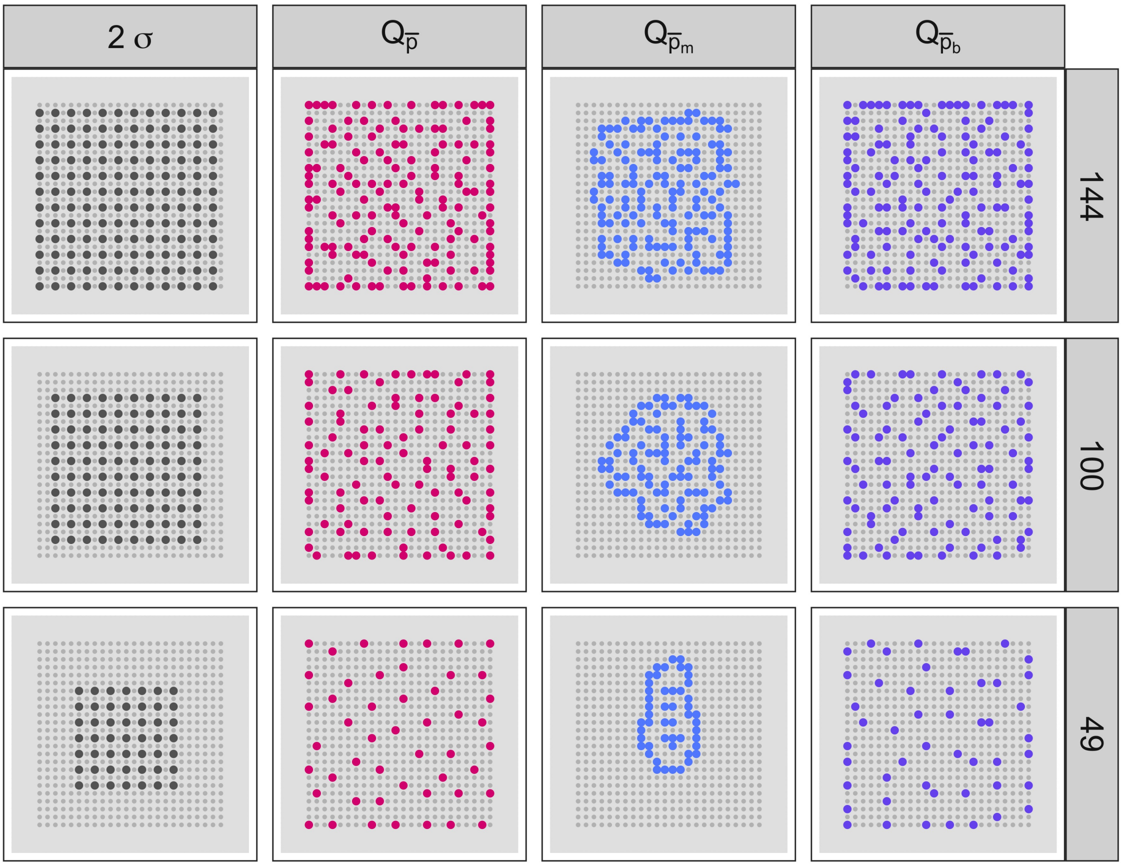


Appendix S2. Figure S1: Full set of designs in the regular study area, including the $2\sigma$ designs as well as the criteria-based designs. From top to bottom, rows represent 144, 100, and 49 traps included in the design, respectively.
