## Appendix S3 for "Optimal sampling design for spatial capture-recapture"

Appendix S3: Vignette of simulation structure. Each row details a single scenario, for which we generated 300 realizations of activity centers, generated detection histories for the specified sampling design, and estimated SCR parameters using two models.

Table S1: Regular study area.

**Regular study area**

| Scenario | Geometry | Effort | Design | Density | Model |
|---|---|---|---|---|---|
| 1 | regular | 144 | $2\sigma$ | uniform | $d.$ |
| 2 | | | | weak | $d.; d_s$ |
| 3 | | | | strong | $d.; d_s$ |
| 4 | | 100 | | uniform | $d.$ |
| 5 | | | | weak | $d.; d_s$ |
| 6 | | | | strong | $d.; d_s$ |
| 7 | | 49 | | uniform | $d.$ |
| 8 | | | | weak | $d.; d_s$ |
| 9 | | | | strong | $d.; d_s$ |
| 10 | | 144 | $Q_{\bar{p}}$ | uniform | $d.$ |
| 11 | | | | weak | $d.; d_s$ |
| 12 | | | | strong | $d.; d_s$ |
| 13 | | 100 | | uniform | $d.$ |
| 14 | | | | weak | $d.; d_s$ |
| 15 | | | | strong | $d.; d_s$ |
| 16 | | 49 | | uniform | $d.$ |
| 17 | | | | weak | $d.; d_s$ |
| 18 | | | | strong | $d.; d_s$ |
| 19 | | 144 | $Q_{\bar{p}_m}$ | uniform | $d.$ |
| 20 | | | | weak | $d.; d_s$ |
| 21 | | | | strong | $d.; d_s$ |
| 22 | | 100 | | uniform | $d.$ |
| 23 | | | | weak | $d.; d_s$ |
| 24 | | | | strong | $d.; d_s$ |
| 25 | | 49 | | uniform | $d.$ |
| 26 | | | | weak | $d.; d_s$ |
| 27 | | | | strong | $d.; d_s$ |
| 28 | | 144 | $Q_{\bar{p}_b}$ | uniform | $d.$ |
| 29 | | | | weak | $d.; d_s$ |
| 30 | | | | strong | $d.; d_s$ |
| 31 | | 100 | | uniform | $d.$ |
| 32 | | | | weak | $d.; d_s$ |
| 33 | | | | strong | $d.; d_s$ |
| 34 | | 49 | | uniform | $d.$ |
| 35 | | | | weak | $d.; d_s$ |
| 36 | | | | strong | $d.; d_s$ |

Table S2: Irregular study area.

**Irregular study area**

| Scenario | Geometry | Effort | Design | Density | Model |
|---|---|---|---|---|---|
| 37 | irregular | 144 | $Q_{\bar{p}}$ | uniform | $d.$ |
| 38 | | | | weak | $d.; d_s$ |
| 39 | | | | strong | $d.; d_s$ |
| 40 | | 100 | | uniform | $d.$ |
| 41 | | | | weak | $d.; d_s$ |
| 42 | | | | strong | $d.; d_s$ |
| 43 | | 49 | | uniform | $d.$ |
| 44 | | | | weak | $d.; d_s$ |
| 45 | | | | strong | $d.; d_s$ |
| 46 | | 144 | $Q_{\bar{p}_m}$ | uniform | $d.$ |
| 47 | | | | weak | $d.; d_s$ |
| 48 | | | | strong | $d.; d_s$ |
| 49 | | 100 | | uniform | $d.$ |
| 50 | | | | weak | $d.; d_s$ |
| 51 | | | | strong | $d.; d_s$ |
| 52 | | 49 | | uniform | $d.$ |
| 53 | | | | weak | $d.; d_s$ |
| 54 | | | | strong | $d.; d_s$ |
| 55 | | 144 | $Q_{\bar{p}_b}$ | uniform | $d.$ |
| 56 | | | | weak | $d.; d_s$ |
| 57 | | | | strong | $d.; d_s$ |
| 58 | | 100 | | uniform | $d.$ |
| 59 | | | | weak | $d.; d_s$ |
| 60 | | | | strong | $d.; d_s$ |
| 61 | | 49 | | uniform | $d.$ |
| 62 | | | | weak | $d.; d_s$ |
| 63 | | | | strong | $d.; d_s$ |
