## Appendix S4 for "Optimal sampling design for spatial capture-recapture"

Appendix S4: Example code implemented in R to generate and plot designs using the optimal sampling design algorithm `scrdesignGA()` for the area of interest in Pakistan. For more information on spatial data preparation and design generation, please see:

https://bookdown.org/chrissuthy/SCR-design-book/

```
library(oSCR)


#----Load data from Pakistan----
data(pakistan)
# This loads the statespace: pakSS
# and the possible trap locations: pakTT


#----Plot this data----
plot(pakSS, asp=1, pch=16, col="grey")
points(pakTT, pch=20, cex = 0.5)


#----Run the design-finding algo----
testdesign <- scrdesignGA(
statespace = pakSS, alltraps = pakTT, # Study area components
ntraps = 25, # Number of available traps
beta0 = 0.2*5,  # Expected data (where beta0 = g0 * k)
sigma = 3, crit = 1,  # Expected data
popsize = 50, keepbest = 5, ngen = 50  # GA settings
)
 # FOR BEST DESIGNS: ngen should be around 1500+
 # See Appendix 1 for a brief discussion of the tuning parameters


#----Plot the results----
par(mfrow=c(1,3))  # Setup plotting area
plot(testdesign, which=4)  # plots all 3 diagnostic plots
par(mfrow=c(1,1))  # Reset plotting area
```
