## Appendix S5 for "Optimal sampling design for spatial capture-recapture"

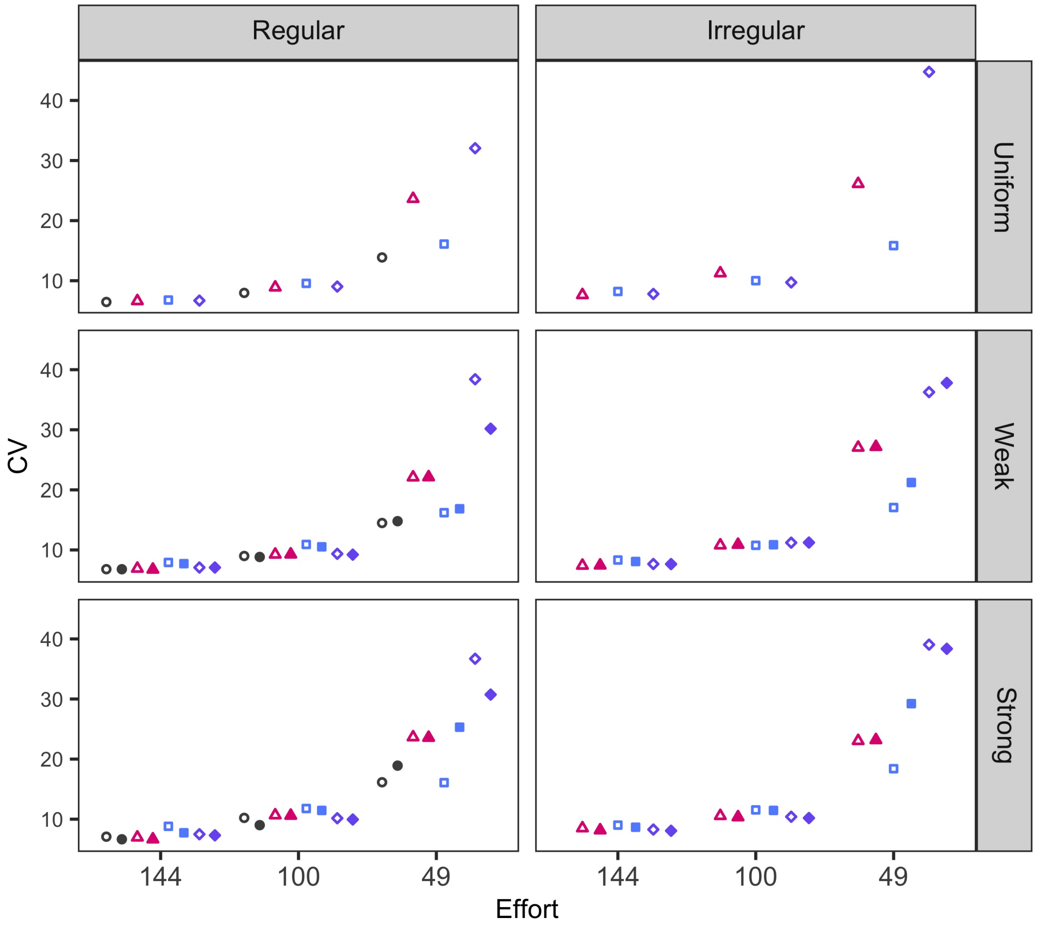


Appendix S5. Figure S1: Coefficient of variation (precision; CV) of estimates of total abundance from the four tested sampling designs under three levels of effort on three density surfaces within two geometries, where estimates are the result of one of two SCR models: density invariant ($d.$, open shapes) or density-varying ($d_s$, closed shapes). The four designs – $2\sigma$, $Q_{\bar{p}}$, $Q_{\bar{p}_m}$, $Q_{\bar{p}_b}$ – are represented by the four shapes: circles, triangles, squares, and diamonds respectively.
