## Appendix S6 for "Optimal sampling design for spatial capture-recapture"

Appendix S6. Table S1: Coefficient of variation (precision; CV) of baseline detection (g0), space use (sigma) and total abundance (EN) for each of the 63 simulation scenarios, in which we varied the design criteria *(Design)*, the shape and accessibility of the landscape *(Geometry, Regular or Irregular)*, the number of traps *(Effort)*, and the underlying density patterns *(Density)*. Results from the null model ($d.$) are reported for all scenarios, and models from the data generating density model ($d_s$) are reported for scenarios with spatially varying density.

| | | | Regular | | | | | | Irregular | | | | | |
|---|---|---|---|---|---|---|---|---|---|---|---|---|---|---|
| | | | $g_0$ | | $\sigma$ | | EN | | $g_0$ | | $\sigma$ | | EN | |
| Effort | Density | Design | $d.$ | $d_s$ | $d.$ | $d_s$ | $d.$ | $d_s$ | $d.$ | $d_s$ | $d.$ | $d_s$ | $d.$ | $d_s$ |
| 49 | uniform | $2\sigma$ | 19.68 | – | 7.28 | – | 13.87 | – | – | – | – | – | – | – |
| | | $Q_{\bar{p}}$ | 22.82 | – | 9.56 | – | 23.64 | – | 25.34 | – | 11.10 | – | 26.12 | – |
| | | $Q_{\bar{p}_m}$ | 16.19 | – | 7.61 | – | 16.10 | – | 18.31 | – | 8.14 | – | 15.84 | – |
| | | $Q_{\bar{p}_b}$ | 22.40 | – | 12.97 | – | 32.05 | – | 22.01 | – | 14.40 | – | 44.76 | – |
| | weak | $2\sigma$ | 18.29 | 18.29 | 7.33 | 7.35 | 14.49 | 14.79 | – | – | – | – | – | – |
| | | $Q_{\bar{p}}$ | 23.05 | 23.05 | 9.86 | 9.87 | 22.09 | 22.12 | 21.94 | 21.94 | 11.52 | 11.45 | 27.04 | 27.17 |
| | | $Q_{\bar{p}_m}$ | 14.91 | 14.91 | 6.48 | 6.49 | 16.20 | 16.85 | 18.80 | 18.80 | 8.05 | 8.04 | 17.06 | 21.22 |
| | | $Q_{\bar{p}_b}$ | 21.20 | 21.20 | 12.59 | 12.20 | 38.41 | 30.18 | 25.28 | 25.28 | 13.85 | 14.01 | 36.26 | 37.80 |
| | strong | $2\sigma$ | 19.24 | 19.24 | 7.62 | 7.61 | 16.15 | 18.91 | – | – | – | – | – | – |
| | | $Q_{\bar{p}}$ | 22.45 | 22.45 | 10.27 | 10.24 | 23.65 | 23.57 | 24.71 | 24.71 | 10.58 | 10.59 | 23.03 | 23.18 |
| | | $Q_{\bar{p}_m}$ | 17.43 | 17.43 | 7.59 | 7.63 | 16.08 | 25.31 | 19.90 | 19.90 | 8.07 | 8.06 | 18.40 | 29.23 |
| | | $Q_{\bar{p}_b}$ | 21.21 | 21.21 | 12.88 | 12.78 | 36.70 | 30.74 | 24.21 | 24.21 | 15.01 | 14.90 | 39.05 | 38.36 |
| 100 | uniform | $2\sigma$ | 12.77 | – | 5.02 | – | 7.97 | – | – | – | – | – | – | – |
| | | $Q_{\bar{p}}$ | 13.02 | – | 6.02 | – | 8.90 | – | 14.50 | – | 6.07 | – | 11.26 | – |
| | | $Q_{\bar{p}_m}$ | 11.02 | – | 4.54 | – | 9.56 | – | 13.00 | – | 5.49 | – | 10.01 | – |
| | | $Q_{\bar{p}_b}$ | 13.74 | – | 5.51 | – | 9.03 | – | 13.74 | – | 5.87 | – | 9.71 | – |
| | weak | $2\sigma$ | 12.72 | 12.72 | 5.16 | 5.16 | 9.00 | 8.83 | – | – | – | – | – | – |
| | | $Q_{\bar{p}}$ | 13.55 | 13.55 | 5.75 | 5.71 | 9.25 | 9.27 | 14.15 | 14.15 | 6.30 | 6.29 | 10.78 | 10.88 |
| | | $Q_{\bar{p}_m}$ | 12.42 | 12.42 | 5.02 | 5.03 | 10.92 | 10.52 | 13.99 | 13.99 | 5.84 | 5.84 | 10.77 | 10.87 |
| | | $Q_{\bar{p}_b}$ | 14.82 | 14.82 | 5.79 | 5.76 | 9.36 | 9.22 | 14.99 | 14.99 | 6.33 | 6.31 | 11.21 | 11.23 |
| | strong | $2\sigma$ | 13.20 | 13.20 | 5.28 | 5.28 | 10.22 | 9.01 | – | – | – | – | – | – |
| | | $Q_{\bar{p}}$ | 14.79 | 14.79 | 6.05 | 6.07 | 10.66 | 10.60 | 14.77 | 14.77 | 6.27 | 6.25 | 10.56 | 10.35 |
| | | $Q_{\bar{p}_m}$ | 10.74 | 10.74 | 4.69 | 4.69 | 11.78 | 11.46 | 12.42 | 12.42 | 5.33 | 5.33 | 11.54 | 11.46 |
| | | $Q_{\bar{p}_b}$ | 14.74 | 14.74 | 5.58 | 5.57 | 10.16 | 9.95 | 15.18 | 15.18 | 6.80 | 6.79 | 10.41 | 10.19 |
| 144 | uniform | $2\sigma$ | 10.86 | – | 4.24 | – | 6.46 | – | – | – | – | – | – | – |
| | | $Q_{\bar{p}}$ | 10.50 | – | 4.14 | – | 6.62 | – | 10.78 | – | 4.70 | – | 7.63 | – |
| | | $Q_{\bar{p}_m}$ | 9.68 | – | 3.95 | – | 6.78 | – | 10.69 | – | 4.20 | – | 8.21 | – |
| | | $Q_{\bar{p}_b}$ | 11.29 | – | 4.43 | – | 6.69 | – | 11.66 | – | 4.74 | – | 7.80 | – |
| | weak | $2\sigma$ | 10.42 | 10.42 | 3.94 | 3.95 | 6.80 | 6.79 | – | – | – | – | – | – |
| | | $Q_{\bar{p}}$ | 11.00 | 11.00 | 4.42 | 4.43 | 6.90 | 6.75 | 11.58 | 11.58 | 4.81 | 4.83 | 7.41 | 7.44 |
| | | $Q_{\bar{p}_m}$ | 8.97 | 8.97 | 3.88 | 3.88 | 7.92 | 7.72 | 10.78 | 10.78 | 4.15 | 4.14 | 8.32 | 8.08 |
| | | $Q_{\bar{p}_b}$ | 10.20 | 10.20 | 4.17 | 4.15 | 7.08 | 7.08 | 11.91 | 11.91 | 4.94 | 4.93 | 7.66 | 7.65 |
| | strong | $2\sigma$ | 10.32 | 10.32 | 4.20 | 4.21 | 7.09 | 6.65 | – | – | – | – | – | – |
| | | $Q_{\bar{p}}$ | 9.83 | 9.83 | 4.46 | 4.45 | 6.98 | 6.66 | 11.23 | 11.23 | 4.58 | 4.56 | 8.51 | 8.17 |
| | | $Q_{\bar{p}_m}$ | 9.47 | 9.47 | 4.04 | 4.03 | 8.81 | 7.74 | 10.65 | 10.65 | 4.54 | 4.54 | 9.01 | 8.66 |
| | | $Q_{\bar{p}_b}$ | 11.56 | 11.56 | 4.34 | 4.32 | 7.50 | 7.31 | 11.97 | 11.97 | 4.48 | 4.49 | 8.28 | 8.07 |
