## Appendix S7 for "Optimal sampling design for spatial capture-recapture"

Appendix S7. Table S1: Scaled root mean square error (accuracy; SRMSE) of baseline detection (g0), space use (sigma) and total abundance (EN) for each of the 63 simulation scenarios, in which we varied the design criteria *(Design)*, the shape and accessibility of the landscape *(, Regular or Irregular)*, the number of traps *(Effort)*, and the underlying density patterns *(Density)*. Results from the null model ($d.$) are reported for all scenarios, and models from the data generating density model ($d_s$) are reported for scenarios with spatially varying density.

| | | | Regular | | | | | | Irregular | | | | | |
|---|---|---|---|---|---|---|---|---|---|---|---|---|---|---|
| | | | $g_0$ | | $\sigma$ | | EN | | $g_0$ | | $\sigma$ | | EN | |
| Effort | Density | Design | $d.$ | $d_s$ | $d.$ | $d_s$ | $d.$ | $d_s$ | $d.$ | $d_s$ | $d.$ | $d_s$ | $d.$ | $d_s$ |
| 49 | uniform | $2\sigma$ | 0.20 | – | 0.07 | – | 0.14 | – | – | – | – | – | – | – |
| | | $Q_{\bar{p}}$ | 0.23 | – | 0.10 | – | 0.26 | – | 0.26 | – | 0.11 | – | 0.29 | – |
| | | $Q_{\bar{p}_m}$ | 0.16 | – | 0.08 | – | 0.16 | – | 0.19 | – | 0.08 | – | 0.16 | – |
| | | $Q_{\bar{p}_b}$ | 0.22 | – | 0.13 | – | 0.39 | – | 0.23 | – | 0.14 | – | 0.56 | – |
| | weak | $2\sigma$ | 0.19 | 0.19 | 0.07 | 0.07 | 0.14 | 0.15 | – | – | – | – | – | – |
| | | $Q_{\bar{p}}$ | 0.23 | 0.23 | 0.10 | 0.10 | 0.24 | 0.24 | 0.22 | 0.22 | 0.11 | 0.11 | 0.31 | 0.31 |
| | | $Q_{\bar{p}_m}$ | 0.15 | 0.15 | 0.06 | 0.06 | 0.16 | 0.17 | 0.19 | 0.19 | 0.08 | 0.08 | 0.17 | 0.22 |
| | | $Q_{\bar{p}_b}$ | 0.21 | 0.21 | 0.12 | 0.12 | 0.47 | 0.38 | 0.25 | 0.25 | 0.14 | 0.14 | 0.43 | 0.45 |
| | strong | $2\sigma$ | 0.20 | 0.20 | 0.08 | 0.08 | 0.17 | 0.20 | – | – | – | – | – | – |
| | | $Q_{\bar{p}}$ | 0.23 | 0.23 | 0.10 | 0.10 | 0.26 | 0.26 | 0.25 | 0.25 | 0.11 | 0.11 | 0.25 | 0.25 |
| | | $Q_{\bar{p}_m}$ | 0.18 | 0.18 | 0.08 | 0.08 | 0.16 | 0.28 | 0.20 | 0.20 | 0.08 | 0.08 | 0.19 | 0.33 |
| | | $Q_{\bar{p}_b}$ | 0.21 | 0.21 | 0.13 | 0.13 | 0.44 | 0.38 | 0.25 | 0.25 | 0.15 | 0.15 | 0.47 | 0.47 |
| 100 | uniform | $2\sigma$ | 0.13 | – | 0.05 | – | 0.08 | – | – | – | – | – | – | – |
| | | $Q_{\bar{p}}$ | 0.14 | – | 0.06 | – | 0.09 | – | 0.15 | – | 0.06 | – | 0.12 | – |
| | | $Q_{\bar{p}_m}$ | 0.11 | – | 0.05 | – | 0.10 | – | 0.13 | – | 0.05 | – | 0.10 | – |
| | | $Q_{\bar{p}_b}$ | 0.14 | – | 0.06 | – | 0.09 | – | 0.14 | – | 0.06 | – | 0.10 | – |
| | weak | $2\sigma$ | 0.13 | 0.13 | 0.05 | 0.05 | 0.09 | 0.09 | – | – | – | – | – | – |
| | | $Q_{\bar{p}}$ | 0.14 | 0.14 | 0.06 | 0.06 | 0.09 | 0.09 | 0.14 | 0.14 | 0.06 | 0.06 | 0.11 | 0.11 |
| | | $Q_{\bar{p}_m}$ | 0.13 | 0.13 | 0.05 | 0.05 | 0.11 | 0.11 | 0.14 | 0.14 | 0.06 | 0.06 | 0.11 | 0.11 |
| | | $Q_{\bar{p}_b}$ | 0.15 | 0.15 | 0.06 | 0.06 | 0.09 | 0.09 | 0.15 | 0.15 | 0.06 | 0.06 | 0.11 | 0.11 |
| | strong | $2\sigma$ | 0.13 | 0.13 | 0.05 | 0.05 | 0.10 | 0.09 | – | – | – | – | – | – |
| | | $Q_{\bar{p}}$ | 0.15 | 0.15 | 0.06 | 0.06 | 0.11 | 0.11 | 0.15 | 0.15 | 0.06 | 0.06 | 0.11 | 0.11 |
| | | $Q_{\bar{p}_m}$ | 0.11 | 0.11 | 0.05 | 0.05 | 0.12 | 0.12 | 0.13 | 0.13 | 0.05 | 0.05 | 0.11 | 0.12 |
| | | $Q_{\bar{p}_b}$ | 0.15 | 0.15 | 0.06 | 0.06 | 0.10 | 0.10 | 0.15 | 0.15 | 0.07 | 0.07 | 0.10 | 0.10 |
| 144 | uniform | $2\sigma$ | 0.11 | – | 0.04 | – | 0.06 | – | – | – | – | – | – | – |
| | | $Q_{\bar{p}}$ | 0.10 | – | 0.04 | – | 0.07 | – | 0.11 | – | 0.05 | – | 0.08 | – |
| | | $Q_{\bar{p}_m}$ | 0.10 | – | 0.04 | – | 0.07 | – | 0.11 | – | 0.04 | – | 0.08 | – |
| | | $Q_{\bar{p}_b}$ | 0.11 | – | 0.04 | – | 0.07 | – | 0.12 | – | 0.05 | – | 0.08 | – |
| | weak | $2\sigma$ | 0.10 | 0.10 | 0.04 | 0.04 | 0.07 | 0.07 | – | – | – | – | – | – |
| | | $Q_{\bar{p}}$ | 0.11 | 0.11 | 0.04 | 0.04 | 0.07 | 0.07 | 0.12 | 0.12 | 0.05 | 0.05 | 0.07 | 0.07 |
| | | $Q_{\bar{p}_m}$ | 0.09 | 0.09 | 0.04 | 0.04 | 0.08 | 0.08 | 0.11 | 0.11 | 0.04 | 0.04 | 0.08 | 0.08 |
| | | $Q_{\bar{p}_b}$ | 0.10 | 0.10 | 0.04 | 0.04 | 0.07 | 0.07 | 0.12 | 0.12 | 0.05 | 0.05 | 0.08 | 0.08 |
| | strong | $2\sigma$ | 0.10 | 0.10 | 0.04 | 0.04 | 0.07 | 0.07 | – | – | – | – | – | – |
| | | $Q_{\bar{p}}$ | 0.10 | 0.10 | 0.04 | 0.04 | 0.07 | 0.07 | 0.11 | 0.11 | 0.05 | 0.05 | 0.09 | 0.08 |
| | | $Q_{\bar{p}_m}$ | 0.09 | 0.09 | 0.04 | 0.04 | 0.09 | 0.08 | 0.11 | 0.11 | 0.05 | 0.05 | 0.09 | 0.09 |
| | | $Q_{\bar{p}_b}$ | 0.12 | 0.12 | 0.04 | 0.04 | 0.07 | 0.07 | 0.12 | 0.12 | 0.04 | 0.04 | 0.08 | 0.08 |
