## Appendix S8 for "Optimal sampling design for spatial capture-recapture"

Appendix S8. Table S1: Percent of simulations failed for each scenario due to a lack of spatial recaptures. For each scenario, we simulated encounter histories until 300 of those were acceptable (i.e., included at least one individual captured in more than one trap) and recorded the number of times that threshold was not met, which we recorded as a 'failure', expressed here as a percentage failed across all simulations within each scenario. Scenarios without failures are not included.

| Scenario | Geometry | Design | Density | Effort | % Failed |
|---|---|---|---|---|---|
| 28 | regular | $Q_{\bar{p}}$ | uniform | 49 | 3.23 |
| 29 | regular | $Q_{\bar{p}}$ | weak | 49 | 3.85 |
| 30 | regular | $Q_{\bar{p}}$ | strong | 49 | 4.46 |
| 34 | regular | $Q_{\bar{p}_b}$ | uniform | 49 | 0.33 |
| 35 | regular | $Q_{\bar{p}_b}$ | weak | 49 | 0.33 |
| 55 | irregular | $Q_{\bar{p}}$ | uniform | 49 | 1.96 |
| 56 | irregular | $Q_{\bar{p}}$ | weak | 49 | 1.32 |
| 57 | irregular | $Q_{\bar{p}}$ | strong | 49 | 1.64 |
