## Supplementary material for "Optimal sampling design for spatial capture-recapture": Metadata S1

Code and data used for the analysis are attached (Data S1) and are publicly available at the author's GitHub page:

`https://github.com/GatesDupont/scr_design_sims`.

The R code, found in the R subdirectory, operates within the file structure of this repository (outline below). Simulations are performed in the `6_sims.r` file, which pulls other files from the directory that are conveniently compiled into the R data file: `workspace/sims_ws.RData`. The simulations file is currently parameterized to run all simulations simultaneously, and for sake of efficiency, the simulations are distributed across several cores.

### Authors

Gates Dupont
University of Massachusetts–Amherst, MA, USA


J. Andrew Royle
U.S. Geological Survey, Laurel, MD, USA


Chris Sutherland
University of Massachusetts–Amherst, MA, USA


| **File** | **Description** |
| --- | --- |
| `R` | Sub-directory containing code for design generations and simulations. |
| `R/0_functions.R` | R code containing functions for simulations, including the simulator() function, which contains the data-generating model. |
| `R/1_SS_regular.R` | R code to generate the regular, square statespace. |
| `R/2_SS_irregular.R` | R code to generate the irregular statespace. |
| `R/3_designs_regular.R` | R code to generate the designs for the regular area. |
| `R/4_designs_irregular.R` | R code to generate the designs for the irregular area. |
| `R/5_sims_gather_ws.R` | R code for gathering data into a single workspace for the simulations. |
| `R/6_sims.R` | R code to run the simulations. Outputs csv file containing rows for each simulation run. |
| `R/7_evaluate` | R code to analyze the results of the simulation. |
| `statespaces` | Sub-directory containing csv files of the statespaces generated in R scripts 1 and 2. |
| `traps` | Sub-directory containing csv files of the possible trapping locations for the regular and irregular statespaces. |
| `designs` | Sub-directory containing csv files of all of the designs evaluated in the simulations. |
| `workspace` | Sub-directory containing an RData file of the workspace compiled in R script 5. |
| `it_out` | Sub-directory that is used to print out each iteration of the simulations. |
| `output` | Sub-directory that is used to print out plots of each simulation as well as the final results file. |
