## Supplementary material for "Optimal sampling design for spatial capture-recapture": Data S1: 0_functions.R

```
# Gates Dupont         #
#    #
# Sept. '19 - July '20 #
# # # # # # # # # # # #

require(oSCR)
require(dplyr)
require(NLMR)
require(raster)
require(viridis)
require(stringr)
require(landscapetools)

select = dplyr::select

"RIGHT"
right = function(text, num_char) {
  substr(text, nchar(text) - (num_char-1), nchar(text))
}


"SIMPLE SS PLOT"
# Plot to check designs
plot_design = function(SS, TT, design){
  plot(SS, asp=1, col="gray80", cex=0.2)
  points(TT, pch=20, col="orange", cex=2)
  points(design, pch=20, col="blue", cex=2.5)
}


"PLANAR GRADIENT"

# Fixed version of nlm_planargradient
r.nlm_planargradient = function (ncol, nrow, resolution = 1, direction = NA, rescale = TRUE) 
{
  checkmate::assert_count(ncol, positive = TRUE)
  checkmate::assert_count(nrow, positive = TRUE)
  checkmate::assert_numeric(direction)
  checkmate::assert_logical(rescale)
  if (is.na(direction)) {
    direction <- stats::runif(1, 0, 360)
  }
  eastness <- sin((pi/180) * direction)
  southness <- cos((pi/180) * direction) * -1
  col_index <- matrix(0:(ncol - 1), nrow, ncol, byrow = TRUE)
  row_index <- matrix(0:(nrow - 1), nrow, ncol, byrow = FALSE)
  gradient_matrix <- (southness * row_index + eastness * col_index)
  gradient_raster <- raster::raster(gradient_matrix)
  raster::extent(gradient_raster) <- c(0, ncol(gradient_raster) * 
                                         resolution, 0, nrow(gradient_raster) * resolution)
  if (rescale == TRUE) {
    gradient_raster <- util_rescale(gradient_raster)
  }
  return(gradient_raster)
}


"DENSIFY"

densify = function(SS, N = 300, landscape = NA, d.beta = 3, seed){
  
  set.seed(seed)
  
  SS = SS[,c("X", "Y")]
  
  # Check parameters
  if(!(landscape %in% c("uniform", "directional", "patchy"))){
    stop("Density must be one of: uniform, directional, patchy")
  }
  
  # Get resolution
  rr = SS %>%
    as.data.frame %>%
    arrange(Y) %>%
    select(X) %>%
    slice(1:2) %>%
    pull(X) %>%
    diff %>% 
    as.numeric
  
  # Calculate number of pixels in each direction (rectangular over the area)
  l.nrow = SS %>% as.data.frame %>% pull(Y) %>% unique %>% sort %>% length
  l.ncol = SS %>% as.data.frame %>% pull(X) %>% unique %>% sort %>% length
  
  # Pull the max n pixels for square
  nside = max(l.ncol, l.nrow)
  
  # Pull the length of the longer side
  lside = max(
    max(SS[,"X"]) - min(SS[,"X"]), 
    max(SS[,"Y"]) - min(SS[,"Y"]))
  
  # Generate the full extent rectangle (might not work for negative axes)
  full = expand.grid(X = seq(min(SS[,"X"]), min(SS[,"X"]) + lside, rr), 
                     Y = seq(min(SS[,"Y"]), min(SS[,"Y"]) + lside, rr))
  
  # Generate densities
  if(landscape == "uniform"){
    surface0 = 1
    surface = 1
  }
  
  if(landscape == "directional"){
    
    # Generate random landscape
    l = nlm_gaussianfield(ncol = nside, nrow = nside, resolution = rr, nug=0,
                          user_seed = seed, rescale = TRUE, 
                          autocorr_range = round(nside))
    
    # Assign the values to the full extent
    full$Z = l@data@values
    
    # Extract raster values from the full extent rectangle to the actual SS and scale
    surface = raster::extract(x = rasterFromXYZ(full), y = as.data.frame(SS))
    
  }
  
  if(landscape == "patchy"){
    
    
    # Simulating the landscape
    l = nlm_gaussianfield(ncol = nside, nrow = nside, resolution = rr, nug=0,
                          user_seed = seed, rescale = TRUE, 
                          autocorr_range = round(nside * 0.06))
    
    # Assign the values to the full extent
    full$Z = l@data@values
    
    # Extract raster values from the full extent rectangle to the actual SS and scale
    surface = raster::extract(x = rasterFromXYZ(full), y = as.data.frame(SS))
  }
  
  # Parameters
  b0 = -1
  b1 = d.beta
  pi = exp(b0 + b1*surface) / sum(exp(b0 + b1*surface)) # Calculating probabilities
  
  # Final SS
  SS = cbind(SS, pi, surface)
  colnames(SS) = c("X", "Y", "density", "surface") 
  
  return(SS)
}


"LOAD ALL FILES"

load_files = function(wd, folder){
  
  dir = paste0(wd, "/", folder)
  
  # Load files
  filenames = list.files(dir, pattern="*.csv", full.names=TRUE)
  objs = lapply(filenames, read.csv)
  
  # Get names
  obj.names = list.files(folder) %>%
    str_remove_all(".csv")
  
  # Assign names
  for(i in 1:length(objs)){
    assign(obj.names[i], objs[[i]], envir = .GlobalEnv)
  }
}


"RMSE CALCULATION"

calc_rmse = function(estimates, parameter){
  diffs = c()
  for(i in 1:length(estimates)){
    diffs[i] = estimates[i] - parameter
  }
  result = sqrt(sum(diffs^2) * (1/length(estimates)))
  
  return(result)
}


"SRMSE CALCULATION"

calc_srmse = function(estimates, parameter){
  diffs = c()
  for(i in 1:length(estimates)){
    diffs[i] = estimates[i] - parameter
  }
  result = (1/parameter) * (sqrt(sum(diffs^2) * (1/length(estimates))))
  
  if(length(result) == 0){
    result = NaN
  } # Handling NA vals
  
  return(result)
}


"SIMULATOR"

#----SIMULATOR----

#Create a simulator function
simulator<- function(traps, ss, N, p0, sigma, K, nsim, it = 1,
                     landscape = NA, d.beta = 3, plot = TRUE, 
                     it.out.dir = NA, plots.out.dir = NA, wd = getwd()) {
  
  # Assign and/or create plotting directory
  plots.out.subdir = paste0(wd, "/", plots.out.dir, "/", "scenario_", it)
  if(!dir.exists(plots.out.subdir)){
    dir.create(plots.out.subdir)
  }
  
  # Initialize data-collection matrix
  simout1 <- matrix(NA, nrow=0, ncol=18)  #c reate empty matrix for output
  colnames(simout1)<- c("p0","sig","d0", "d.beta", # estimates
                        "nind", # number of individuals (length of first dimmension of y)
                        "nind.c", "nind.r", "r_s", "avg.dets", "avg.r", "avg.nlocs", "avg.nspatcaps", # manually calculated summary stats
                        "avg.caps","avg.spatial","mmdm", # summary stats from oSCR (but see metric definitions)
                        "failures","EN","oSCR_model") # other components
  
  # Initialize while loop starting values
  sim = 1
  sim_try = 0
  total_its = 0
  
  # Get nsim acceptable simulations
  while(sim < (nsim + 1)){
    
    # Update loop
    total_its = total_its + 1
    
    # Tell the user what's going on:
    print(paste("Simulation Number", sim_try + 1, sep = " ")) # keep track
    cat("size of state-space: ", nrow(ss), " pixels", fill=TRUE)
    cat(paste0("\n Try ", sim_try + 1, "\n"))
    
    # Adding density surface to statespace
    statespace = densify(SS = ss, landscape = landscape, d.beta = d.beta, seed = sim)
    
    # Sset seed after internal use of set.seed() in densify()
    seed = total_its
    set.seed(seed)
    
    # Sampling activity centers
    ac = as.numeric()
    for(i in 1:N){
      ac[i] = base::sample(x = nrow(statespace), size = 1, prob = statespace[,"density"])
    }
    s = statespace[ac, c("X", "Y") ]
    
    # Make the state space data frame
    myss <- as.data.frame(statespace)[,c("X", "Y", "surface")]
    myss$Tr <- 1
    myss <- list(myss)
    class(myss) <- "ssDF"
    
    # individual-trap distance matrix
    D <- e2dist(s,traps)
    
    # Compute detection probabilities:
    pmat <- p0*exp(-D*D/(2*sigma*sigma)) # p for all inds-traps p_ij
    ntraps <- nrow(traps)
    y <- array(0, dim=c(N, ntraps, K)) # empty 3D array (inds by traps by occ)
    
    for(i in 1:N){# loop through each individual/activity center
      for(j in 1:ntraps){# loop through each trap
        y[i,j,1:K]<- rbinom(K, 1, pmat[i,j]) # y ~ binomial(p_ijk)
      }
    }
  
    ncap <- apply(y,c(1), sum)       # sum of captures for each individual
    y.all = y                        # for summary stats
    y <- y[ncap>0,,]                 # reduce the y array to include only captured individuals
    
    # Some summary information, that is actually printed for you later with "print(scrFrame)"
    caps.per.ind.trap <- apply(y,c(1,2),sum) #shows # capts for each indv across all traps
    
    # Check for captures
    check.y = length(dim(y)) %>%
      if(. > 2){return(TRUE)} else {return(FALSE)}
    
    # Check for spatial recaps
    check.sp_recaps = as.matrix((caps.per.ind.trap > 0) + 0) %>%
      rowSums() %>%
      c(.,-1) %>% # This is just to avoid warning messages due to empty lists
      max %>%
      if(. > 1){return(TRUE)} else {return(FALSE)}
    
    check = 0 # Clear from previous iteration
    check = check.y + check.sp_recaps
    
    # Checking for sp.recaps implies getting caps, 
    # but for troubleshooting good to keep both checks
    
    if(check != 2){
      
      #plot(rasterFromXYZ(statespace[,c(1,2,4)]), col = rev(viridis(1000))) # plot the state space
      #points(s, pch = 20, col = "white")
      #points(s)
      
      #simout1 = rbind(simout1, rep(NA, ncol(simout1))) # Turn this off for the big run
      
    } else if(check ==2){
      
      # Make the SCRframe
      colnames(traps)<- c("X","Y")
      sf <- make.scrFrame(caphist=list(y), traps=list(traps))
      
      # Plotting
      if(plot == TRUE){
        
        # Create individual plot output file
        plots.out.dir.file = paste0(plots.out.subdir, "/", "sim_", it, "_", sim, ".png")
        png(filename = plots.out.dir.file, width = 500, height = 500)
        
        # Make plot
        plot(rasterFromXYZ(statespace[,c(1,2,4)]), col = rev(viridis(1000))) # plot the state space
        points(s, pch = 20, col = "white")
        points(s)
        spiderplot(sf, add=TRUE)
        dev.off()
        
      }
      
      
      #----SCR SUMMARY STATS----
      # Collapse occasion dim
      collap.y.tot = apply(y.all, 1:2, sum)
      
      # Copy (for later)
      collap.y.sum = collap.y.tot
      
      # Number of captures per individual
      ncaps.per.ind = rowSums(collap.y.tot)
      
      # Convert this to binary and count ntraps per individual
      collap.y.sum[collap.y.sum > 0] = 1
      ntraps.per.ind = rowSums(collap.y.sum)
      ntraps.per.capInd = ntraps.per.ind[ntraps.per.ind>0]
      
      # Basics, n and r
      nind.c = length(ncaps.per.ind[ncaps.per.ind>0])        # number of individuals with captures
      nind.r = length(ncaps.per.ind[ncaps.per.ind>1])        # number of individuals with recaptures
      
      # For only captured/detected individuals...
      r_s = sum(ntraps.per.capInd - 1)                  # total sp recaps... sum (number of traps per detected ind - 1)
      avg.dets = mean(ncaps.per.ind[ncaps.per.ind>0])   # avg. of the number of caps per detected individual
      avg.r = mean(ntraps.per.capInd - 1)               # avg  of the number of recaps per detected ind
      avg.nlocs = mean(ntraps.per.capInd)               # avg of the number of unique locations per individual
      avg.nspatcaps = mean(ntraps.per.capInd - 1)       # avg of the number of spatial captures per individual
      
      # Finally summary stats object
      SCR_summary_stats = list(nind.c, nind.r, r_s, avg.dets, avg.r, avg.nlocs, avg.nspatcaps)
      names(SCR_summary_stats) = c("nind.c", "nind.r", "r_s", "avg.dets", "avg.r", "avg.nlocs", "avg.nspatcaps")
      
      
      #----Continuing to model fitting----
      # Fit a basic model SCR0
      out1 <- oSCR.fit(model=list(D~1,p0~1,sig~1), scrFrame = sf, ssDF=myss, trimS = 4*sigma)
      
      # UNIFORM DENSITY Estimates: d_.
      stats <- print(sf)[[1]]  # pulls avg caps, avg spatial caps, and mmdm
      est <- out1$outStats$mle       # pulls p0, sigma, and d0 estimates from the model
      en = get.real(out1, type="dens", newdata=data.frame(session=factor(1)),
                    d.factor=nrow(out1$ssDF[[1]]))[1,1] # Total abundance
      
      # Append to data-collection matrix
      sim_vals = c(plogis(est[1]), exp(est[2]), exp(est[3]), NA, dim(y)[1], SCR_summary_stats, stats, sim_try, en, 1)
      simout1 = rbind(simout1, sim_vals)
      
      
      # INHOMOGENOUS DENSITY Estimate: d_s
      if(landscape != "uniform"){
        
        out2 <- oSCR.fit(model=list(D~surface,p0~1,sig~1), scrFrame = sf, ssDF=myss, trimS = 4*sigma)
        est2 = out2$outStats$mle
        en2 = sum(get.real(out2, type="dens")[[1]]$estimate) 
        sim_vals2 = c(plogis(est[1]), exp(est2[2]), exp(est2[3]), est2[4], dim(y)[1], SCR_summary_stats, stats, sim_try, en2, 2)
        simout1 = rbind(simout1, sim_vals2)
        
      } 
      
    }
    
    
    if(!is.na(it.out.dir) & is.character(it.out.dir)){
      it.out.dir.file = paste0(wd, "/", it.out.dir, "/", "sim", it, ".txt")
      write.table(simout1, file = it.out.dir.file)
    } 
    
    
    # Updating while() loop
    if(check != 2){
      sim_try <- sim_try + 1
    } else {
      sim <- sim + 1
      sim_try = 0
    }
    
    
  }
  
  return(simout1)
  
}
```
