## Supplementary material for "Optimal sampling design for spatial capture-recapture": Data S1: 1_SS_regular.R

```
# Gates Dupont         #
#    #
# Sept. '19 - July '20 #
# # # # # # # # # # # #

library(dplyr)
library(NLMR)
library(raster)
library(viridis)
source("R/0_functions.R")
select = dplyr::select
wd = getwd()

#----Primary parameter----

sig = 0.8


#----Statespace----

# Define the state-space
ss.xlim <- ss.ylim <- c(0, 24)

# Make the state space
rr <- 0.5
SS <- as.matrix(expand.grid(X = seq(ss.xlim[1]+rr/2, ss.xlim[2]-rr/2,rr),
                            Y = seq(ss.ylim[1]+rr/2, ss.ylim[2]-rr/2,rr)))

#----Traps 2----

# Make the traps
rr <- sig
tr.xlim <- tr.ylim <- c(3*sig, (ss.xlim[2])-(3*sig))
TT <- as.matrix(expand.grid(X = seq(tr.xlim[1]+rr/2, tr.xlim[2]-rr/2,rr),
                            Y = seq(tr.ylim[1]+rr/2, tr.ylim[2]-rr/2,rr)))


"CHECK"

#----Plottting statespaces and traps----

plot(SS, pch=20, col="gray80", cex=0.1, asp=1)
points(TT, pch=20)


#----Writing SS csv s----

# STATESPACES
dir = paste0(wd, "/statespaces")

if(!dir.exists(dir)){
  dir.create(dir)
}

file = paste0(dir, "/SS_regular.csv")
write.csv(SS, file, row.names = FALSE)

# TRAPS
dir = paste0(wd, "/traps")

if(!dir.exists(dir)){
  dir.create(dir)
}

file = paste0(dir, "/possible_traps_regular", ".csv")
write.csv(TT, file, row.names = FALSE)
```
