## Supplementary material for "Optimal sampling design for spatial capture-recapture": Data S1: 2_SS_irregular.R

```
# Gates Dupont         #
#    #
# Sept. '19 - July '20 #
# # # # # # # # # # # #

# Tidyverse
library(dplyr)
library(purrr)
library(tidyr)
library(reshape2)
library(ggplot2)
library(ggpubr)
library(ggforce)
library(viridis)

# Spatial
library(raster)
library(rgdal)
library(rgeos)
library(NLMR)
library(landscapetools)
library(grid)
library(gdistance)
library(sf)
library(plotrix)

# Source the functions
source("R/0_functions.R")

# Namespace
extract = raster::extract
group_by = dplyr::group_by
select = dplyr::select

# Working directory for r proj
wd = getwd()

"SETTING UP IRREGULAR SHAPE"

#----Loading spatial data----
grids <- readOGR("data/pakistan/spatial_data/5x5km.shp")
locs <- readOGR("data/pakistan/spatial_data/Astore_Camera_Trap_New.shp")
cams <- readOGR("data/pakistan/spatial_data/Camera_TL.shp")
cams2 <- readOGR("data/pakistan/spatial_data/Camera_TL_Refined.shp")
roads <- readOGR("data/pakistan/spatial_data/Roads.shp")
vills <- readOGR("data/pakistan/spatial_data/Villages.shp")
excluded <- readOGR("data/pakistan/spatial_data/Excluded_areas.shp")
dem1 <- raster("data/pakistan/spatial_data/DEM_NAs_UTM_100m.tif")
studyarea <- readOGR("data/pakistan/spatial_data/Astore_Camera_Trap_New.shp")
studyarea <- readOGR("data/pakistan/spatial_data/Study_Whole_New.shp")


#----IRREGULAR STATESPACE----

# Parameters from reg_big_ss
reg_ss.side = 24
reg_ss.sig = 0.8
reg_ss.rr = 0.5
reg_ss.px = 2304
reg_ss.N = 300

# Length of side is 3 sig on both ends, squared, divided by npix
areaofspace = ((reg_ss.side - (2*3*reg_ss.sig))^2)/(reg_ss.rr^2)
areaoftotal = (reg_ss.side^2)/(reg_ss.rr^2)
sigmaratio = reg_ss.sig/sqrt(reg_ss.rr^2)

# Loading pakistan data
pak = st_read("data/pakistan/spatial_data", "Study_Whole_New")
pakpoints = st_sample(pak, size = areaofspace, type = "regular", exact = TRUE) 

# Making irregular SS
irreg_rr = as_Spatial(pakpoints)@coords %>%
  as.data.frame %>%
  pull(coords.x1) %>%
  sort() %>%
  diff %>%
  max

# Add buffer to study area
irreg.sigma.x1000 = sigmaratio*irreg_rr 
pak_wBuff = st_buffer(x = pak, dist = 3*irreg.sigma.x1000)

# Generate area pixels & ensure equal area btwn reg and irreg
check_area = 0 
while (check_area != areaoftotal) {
  pakpoints_wBuff = st_sample(pak_wBuff, size = areaoftotal, type = "regular", exact = TRUE) 
  check_area = length(pakpoints_wBuff)
}

# Create irreg ss
SS = cbind(1:length(pakpoints_wBuff),st_coordinates(pakpoints_wBuff)/1000)[,2:3]


"GENERATE TRAPS"

# Namespace for sigma
sigma.x1000 = irreg.sigma.x1000


#----Analayze data to generate possible traps----

# Spatial data
elevation.sub <- crop(dem1, extent(pak_wBuff))
elev <- mask(elevation.sub, pak_wBuff)
slop <- terrain(elev, opt='slope', unit='degrees', neighbors=8)
tri <- terrain(elev, opt='TRI', neighbors=8)
nogo <- rasterize(excluded,elev)

# Making possible camera locs
ext = as_Spatial(pak) %>% extent() %>% as.matrix()
all.locs = expand.grid(X = seq(ext["x","min"], ext["x","max"], by=sigma.x1000),
                       Y = seq(ext["y","min"], ext["y","max"], by=sigma.x1000)) %>%
  SpatialPoints(proj4string = crs(elev)) %>%
  .[as_Spatial(pak),]

# Filter data for potential trap locations
mytraps = data.frame(X=coordinates(all.locs)[,1],
                     Y=coordinates(all.locs)[,2],
                     elev = extract(elev,coordinates(all.locs)),
                     slope = extract(slop,coordinates(all.locs)),
                     TRI = extract(tri,coordinates(all.locs)),
                     access = ifelse(is.na(extract(nogo,coordinates(all.locs))),1,0)) %>%
  filter(access == 1) %>%
  filter(slope <= 45) %>%
  filter(elev <= 4500) %>% # use 4200
  select(X, Y) %>%
  mutate(X=X/1000, Y=Y/1000) %>%
  as.matrix()

# Plotting possible trap locations
plot(SS,pch=16, asp=1,cex=0.3, col="gray80")
points(mytraps,pch=16,col="orange",cex=0.6)


"OUTPUT"

#----Writting files----

# STATESPACES
dir = paste0(wd, "/statespaces")

if(!dir.exists(dir)){
  dir.create(dir)
}

file = paste0(dir, "/SS_irregular.csv")
write.csv(SS, file, row.names = FALSE)


# TRAPS
dir = paste0(wd, "/traps")

if(!dir.exists(dir)){
  dir.create(dir)
}

file = paste0(dir, "/possible_traps_irregular.csv")
write.csv(mytraps, file, row.names = FALSE)
```
