## Supplementary material for "Optimal sampling design for spatial capture-recapture": Data S1: 3_designs_regular.R

```
"HEY! LOOK! SETUP"

# Parallel
ncores = 19 # To parallelize design finding

# GA
ngen = 1500   # This should be >1500 for smooth designs
popsize = 200 # Default parameters in GA
keepbest = 20

# SCR Parameters
p0 = 0.2
K = 5
sig = 0.8

# # # # # # # # # # # # # # # # # # # # # # # # # # # # # # # # # # # # # # # # # # # # # # # 
# Gates Dupont         #
#    #
# Sept. '19 - July '20 #
# # # # # # # # # # # #


library(oSCR)
library(doParallel)
library(kofnGA)

# Source the functions
source("R/0_functions.R")

# Working directory
wd = getwd()


#----Load data----
file = paste0(wd, "/statespaces/SS_regular.csv")
SS = read.csv(file)

file = paste0(wd, "/traps/possible_traps_regular.csv")
TT = read.csv(file)


#---Number of traps for 2sigma-----
ntrapsA = 12^2 # full coverage
ntrapsB = 10^2
ntrapsC = 7^2


#----Manually generating 2 sigma grid----

# Full coverage
designA.2sig = expand.grid(X = seq(min(TT[,1]), max(TT[,1]), by=2*sig)[1:sqrt(ntrapsA)],
                          Y = seq(min(TT[,1]), max(TT[,1]), by=2*sig)[1:sqrt(ntrapsA)])

# Roughly a two thirds coverage
designB.2sig = expand.grid(X = seq(min(TT[,1]), max(TT[,1]), by=2*sig)[1:sqrt(ntrapsB)],
                          Y = seq(min(TT[,1]), max(TT[,1]), by=2*sig)[1:sqrt(ntrapsB)]) %>%
  mutate(X = X + (2*sig), Y = Y + (2*sig)) 

# Roughly one thirds coverage
designC.2sig = expand.grid(X = seq(min(TT[,1]), max(TT[,1]), by=2*sig)[1:sqrt(ntrapsC)],
                          Y = seq(min(TT[,1]), max(TT[,1]), by=2*sig)[1:sqrt(ntrapsC)]) %>%
  mutate(X = X + (5*sig), Y = Y + (5*sig)) 


#----Checking configurations----

par(mfrow = c(1,3))

plot_design(SS, TT, designA.2sig)
plot_design(SS, TT, designB.2sig)
plot_design(SS, TT, designC.2sig)

par(mfrow = c(1,1))


#----GA designs----

"A"

# pbar
cl = makeCluster(ncores)
clusterExport(cl, varlist = c("e2dist"), envir = environment())
designA.pbar.GA =
  scrdesignGA(
    statespace = SS, alltraps = TT, ntraps = ntrapsA, # Study area
    sigma = sig, beta0 = log(p0*K), crit = 1, # SCR parameters
    popsize = popsize, keepbest = keepbest, ngen = ngen, # GA parameters
    cluster = cl
  )
stopCluster(cl)
designA.pbar = designA.pbar.GA$optimaltraps

# p2bar
cl = makeCluster(ncores)
clusterExport(cl, varlist = c("e2dist"), envir = environment())
designA.p2bar.GA =
  scrdesignGA(
    statespace = SS, alltraps = TT, ntraps = ntrapsA, # Study area
    sigma = sig, beta0 = log(p0*K), crit = 2, # SCR parameters
    popsize = popsize, keepbest = keepbest, ngen = ngen, # GA parameters
    cluster = cl
  )
stopCluster(cl)
designA.p2bar = designA.p2bar.GA$optimaltraps

# pcombo
cl = makeCluster(ncores)
clusterExport(cl, varlist = c("e2dist"), envir = environment())
designA.pcombo.GA =
  scrdesignGA(
    statespace = SS, alltraps = TT, ntraps = ntrapsA, # Study area
    sigma = sig, beta0 = log(p0*K), crit = 3, # SCR parameters
    popsize = popsize, keepbest = keepbest, ngen = ngen, # GA parameters
    cluster = cl
  )
stopCluster(cl)
designA.pcombo = designA.pcombo.GA$optimaltraps


"B"

# pbar
cl = makeCluster(ncores)
clusterExport(cl, varlist = c("e2dist"), envir = environment())
designB.pbar.GA =
  scrdesignGA(
    statespace = SS, alltraps = TT, ntraps = ntrapsB, # Study area
    sigma = sig, beta0 = log(p0*K), crit = 1, # SCR parameters
    popsize = popsize, keepbest = keepbest, ngen = ngen, # GA parameters
    cluster = cl
  )
stopCluster(cl)
designB.pbar = designB.pbar.GA$optimaltraps

# p2bar
cl = makeCluster(ncores)
clusterExport(cl, varlist = c("e2dist"), envir = environment())
designB.p2bar.GA =
  scrdesignGA(
    statespace = SS, alltraps = TT, ntraps = ntrapsB, # Study area
    sigma = sig, beta0 = log(p0*K), crit = 2, # SCR parameters
    popsize = popsize, keepbest = keepbest, ngen = ngen, # GA parameters
    cluster = cl
  )
stopCluster(cl)
designB.p2bar = designB.p2bar.GA$optimaltraps

# pcombo
cl = makeCluster(ncores)
clusterExport(cl, varlist = c("e2dist"), envir = environment())
designB.pcombo.GA =
  scrdesignGA(
    statespace = SS, alltraps = TT, ntraps = ntrapsB, # Study area
    sigma = sig, beta0 = log(p0*K), crit = 3, # SCR parameters
    popsize = popsize, keepbest = keepbest, ngen = ngen, # GA parameters
    cluster = cl
  )
stopCluster(cl)
designB.pcombo = designB.pcombo.GA$optimaltraps


"C"

# pbar
cl = makeCluster(ncores)
clusterExport(cl, varlist = c("e2dist"), envir = environment())
designC.pbar.GA =
  scrdesignGA(
    statespace = SS, alltraps = TT, ntraps = ntrapsC, # Study area
    sigma = sig, beta0 = log(p0*K), crit = 1, # SCR parameters
    popsize = popsize, keepbest = keepbest, ngen = ngen, # GA parameters
    cluster = cl
  )
stopCluster(cl)
designC.pbar = designC.pbar.GA$optimaltraps

# p2bar
cl = makeCluster(ncores)
clusterExport(cl, varlist = c("e2dist"), envir = environment())
designC.p2bar.GA =
  scrdesignGA(
    statespace = SS, alltraps = TT, ntraps = ntrapsC, # Study area
    sigma = sig, beta0 = log(p0*K), crit = 2, # SCR parameters
    popsize = popsize, keepbest = keepbest, ngen = ngen, # GA parameters
    cluster = cl
  )
stopCluster(cl)
designC.p2bar = designC.p2bar.GA$optimaltraps

# pcombo
cl = makeCluster(ncores)
clusterExport(cl, varlist = c("e2dist"), envir = environment())
designC.pcombo.GA =
  scrdesignGA(
    statespace = SS, alltraps = TT, ntraps = ntrapsC, # Study area
    sigma = sig, beta0 = log(p0*K), crit = 3, # SCR parameters
    popsize = popsize, keepbest = keepbest, ngen = ngen, # GA parameters
    cluster = cl
  )
stopCluster(cl)
designC.pcombo = designC.pcombo.GA$optimaltraps


#----Write out the files----
designs = list(designA.2sig, designA.pbar, designA.p2bar, designA.pcombo,
               designB.2sig, designB.pbar, designB.p2bar, designB.pcombo,
               designC.2sig, designC.pbar, designC.p2bar, designC.pcombo)

filename_ntraps = rep(c("designA_", "designB_", "designC_"), each = 4)
filename_design = rep(c("2sig_", "pbar_", "p2bar_", "pcombo_"), 3)

dir = paste0(wd, "/designs")
if(!dir.exists(dir)){
  dir.create(dir)
}

for (i in 1:12) {
  file = paste0(dir, "/", filename_ntraps[i], filename_design[i], "_regular.csv")
  write.csv(designs[[i]], file, row.names = FALSE)
}
```
