## Supplementary material for "Optimal sampling design for spatial capture-recapture": Data S1: 5_sims_gather_ws.R

```
# Gates Dupont         #
#    #
# Sept. '19 - July '20 #
# # # # # # # # # # # #

library(stringr)

source("R/0_functions.R")

wd = getwd()

"LOADING DATA"

#----Load all data----

load_files(wd, folder = "statespaces")
load_files(wd, folder = "traps")
load_files(wd, folder = "designs")


#----Organizing----

statespaces = mget(c(rep("SS_regular", 36), rep("SS_irregular", 63-36)))
designs = mget(c(
  # REGULAR
  "designA_2sig__regular", 
  "designA_pbar__regular", "designA_p2bar__regular", "designA_pcombo__regular",
                 
  "designB_2sig__regular", 
  "designB_pbar__regular", "designB_p2bar__regular", "designB_pcombo__regular",
                 
  "designC_2sig__regular", 
  "designC_pbar__regular", "designC_p2bar__regular", "designC_pcombo__regular",
                 
  # "Irregular"
  "designA_pbar__irregular", "designA_p2bar__irregular", "designA_pcombo__irregular",
                 
  "designB_pbar__irregular", "designB_p2bar__irregular", "designB_pcombo__irregular",
                 
  "designC_pbar__irregular", "designC_p2bar__irregular", "designC_pcombo__irregular"
  )) %>%
  rep(each = 3)


#----Formatting----

reg_ss.sig = possible_traps_regular %>% as.data.frame %>% 
  arrange(Y) %>% slice(1:2) %>% pull(X) %>% diff

irreg_ss.sig = possible_traps_irregular %>% as.data.frame %>% 
  arrange(Y) %>% slice(1:2) %>% pull(X) %>% diff

# Sim parameters
sig_true = c(rep(reg_ss.sig, 36), rep(irreg_ss.sig, 63-36))
N = rep(300, 63)
p0_true = rep(0.2, 63)
K = rep(5, 63)
d0_true = c(rep(300/nrow(SS_regular), 36), rep(300/nrow(SS_irregular), 63-36))


# Descriptives
names_design = c(
  rep(rep(c("2sigma", "pbar", "p2bar", "pcombo"), each = 3), 3),
  rep(rep(c("pbar", "p2bar", "pcombo"), each = 3), 3))

names_coverage = c(
  rep(c("A", "B", "C"), each = 12),
  rep(c("A", "B", "C"), each = 9))

names_geom = c(rep("regular", 36), rep("irregular", 63-36))

names_density = rep(c("uniform", "patchy", "directional"), 21)

# Final structure
sim_structure = data.frame(
  i = as.character(1:63),
  names_geom = names_geom,
  names_coverage = names_coverage,
  names_design = names_design,
  names_density,
  statespaces = names(statespaces),
  designs = names(designs),
  sig_true = sig_true,
  N = N,
  p0_true = p0_true,
  K = K,
  d0_true = d0_true
)
View(sim_structure)


#----Save the essentials----
keep = c("wd", "names_geom", "names_coverage", "names_design",
         "names_density", "statespaces", "designs", "sig_true", "N",
         "p0_true", "K", "d0_true")

rm(list = setdiff(ls(), keep))


#----Saving as RData workspace----

dir = paste0(wd, "/", "workspace")

if(!dir.exists(dir)){
  dir.create(dir)
}

file = paste0(dir, "/sims_ws.RData")
save.image(file)
```
