## Supplementary material for "Optimal sampling design for spatial capture-recapture": Data S1: 6_sims.R

```
"HEY! LOOK! Setup"

ncores = 15
nsim = 300 # This should be 300
wd = getwd()
output.dir = "output"
it.out.dir = "it_out"
plots.out.dir = "output/sim_plots"
d.beta_true = 1.2


if(!dir.exists(output.dir)){
  dir.create(output.dir)
}

if(!dir.exists(it.out.dir)){
  dir.create(it.out.dir)
}

if(!dir.exists(plots.out.dir)){
  dir.create(plots.out.dir)
}


# # # # # # # # # # # # # # # # # # # # # # # # # # # # # # # # # # # # # # # # # # # # # # # #
# Gates Dupont         #
#    #
# Sept. '19 - July '20 #
# # # # # # # # # # # #

library(oSCR)
library(dplyr)
library(raster)
library(foreach)
library(doParallel)
library(viridis)
library(landscapetools)

extract = raster::extract
group_by = dplyr::group_by
select = dplyr::select

source("R/0_functions.R")

file = "workspace/sims_ws.RData"
load(file)


#----Running all simulations----

set.seed(1) # for repro

# Making parallel cluster
cl = makeCluster(ncores)
registerDoParallel(cl)

simout = list()

t0 = Sys.time()
simout = foreach(i=1:length(designs), .packages=c(.packages()), .errorhandling = "pass") %dopar% {
  
  # For repro
  set.seed(i)
  
  # Create first row
  simout_i = simulator(traps = designs[[i]], ss = statespaces[[i]], 
                       N = N[i], p0 = p0_true[i], sigma = sig_true[i], K = K[i],
                       landscape = names_density[i], d.beta = d.beta_true, wd = getwd(),
                       it.out.dir = it.out.dir, plots.out.dir = plots.out.dir,
                       it = as.character(i), nsim = nsim, plot = TRUE) %>%
    apply(., 2, as.numeric) %>%
    as.data.frame() %>%
    mutate(geom = names_geom[i]) %>%
    mutate(coverage = names_coverage[i]) %>%
    mutate(design = names_design[i]) %>%
    mutate(density = names_density[i]) %>%
    # True values
    mutate(N_true = N[i],
           p0_true = p0_true[i],
           sig_true = sig_true[i],
           d0_true = d0_true[i],
           d.beta_true = d.beta_true,
           i = i) %>%
    select(i, everything())
  
  write.table(simout_i, paste0(it.out.dir, "/", "all_sims_scenario_", as.character(i), ".txt"))
  
  return(simout_i)
}
stopCluster(cl)
tf = Sys.time()

if(length(simout) > 1){
  print(tf-t0)
} else {
  stop("Simulations did not run")
}

#----Writing results----
results = do.call(rbind, simout)

# Write sims output
file = paste0(output.dir, "/", "results.csv")
write.csv(results, file, row.names = FALSE)

# Write total time
file = paste0(output.dir, "/", "stopwatch.csv")
stopwatch = data.frame(time_start = t0, time_stop = tf)
write.csv(stopwatch, file)
```
