## Supplementary material for "Optimal sampling design for spatial capture-recapture": Data S1: 7_evaluate.R

```
wd = getwd()

# Gates Dupont         #
#    #
# Sept. '19 - July '20 #
# # # # # # # # # # # #


library(Hmisc)
library(dplyr)
library(tidyr)
library(forcats)
library(viridis)
select =  dplyr::select
ibm = c("#648FFF", "#785EF0", "#DC267F", "#FE6100", "#FFB000")

file = paste0(wd, "/R/", "0_functions.R")
source(file)


"_____________________Formatting Data_____________________"

#----Load data----
file <- paste0(wd, "/data/", "results_061320.csv")
df0 <- read.csv(file)


#----Ordering factors---
df <- df0 %>%
  mutate(X = 1:nrow(df0)) %>%
  select(i, X, everything()) %>%
  mutate(oSCR_model = replace(oSCR_model, oSCR_model == 1,  "d.")) %>%
  mutate(oSCR_model = replace(oSCR_model, oSCR_model == 2,  "ds")) %>%
  mutate(geom = factor(geom, levels = c("regular", "irregular"))) %>%
  mutate(density = factor(density, levels = c("uniform", "patchy", "directional"))) %>%
  mutate(density = fct_recode(density,  "uniform" = "uniform", "weak" = "patchy", "strong" = "directional" )) %>%
  arrange(X)

fail_df <- df %>%
  filter(oSCR_model != "ds") %>%
  select(i, geom, design, density, coverage, failures) %>%
  group_by(i, geom, design, density, coverage) %>%
  summarise(mean_n_fails = 100*((sum(failures))/(300+sum(failures))) ) %>%
  filter(mean_n_fails > 0)


#----Go long!----
out <- df %>%
  gather(Parameter, Value, c(p0:d.beta, EN), factor_key=TRUE) %>%
  mutate(Truth = ifelse(Parameter == "d0" & geom == "regular", unique(d0_true)[1],
                        ifelse(Parameter == "d0" & geom == "irregular", unique(d0_true)[2],
                               ifelse(Parameter == "p0", p0_true,
                                      ifelse(Parameter == "sig" & geom == "regular", unique(sig_true)[1],
                                             ifelse(Parameter == "sig" & geom == "irregular", unique(sig_true)[2],
                                                    ifelse(Parameter == "EN", N_true,
                                                           ifelse(Parameter == "d.beta" & oSCR_model == "ds", d.beta_true, NA))))))))


"_____________________Summary_____________________"

#----Summary----
d.sum <- df %>%
  filter(oSCR_model != "ds") %>%
  select(i, X, geom, density, design, coverage, nind, avg.caps, avg.spatial) %>%
  group_by(geom, density, design, coverage) %>%
  mutate(
    avg.nind = mean(nind, na.rm = T),
    total.avg.caps = mean(avg.caps, na.rm = T),
    total.avg.spatial = mean(avg.spatial, na.rm = T),
    i = unique(i)
  ) %>%
  select(i, geom, density, design, coverage, avg.nind, total.avg.caps, total.avg.spatial)  %>%
  unique() %>%
  mutate_if(is.numeric, round, 2) %>%
  arrange(i)


"_____________________Evaluations_____________________"

#----Evaluations----
evals <- out %>%
  mutate(SME = 100 * (Value-Truth)/Truth) %>%
  group_by(geom, density, design, coverage, oSCR_model, Parameter) %>%
  mutate(CV = 100 * (sd(na.omit(Value)))/mean(Value, na.rm = T),
         SRMSE = calc_srmse(na.omit(Value), na.omit(unique(Truth)))) %>%
  summarise(SME_L = quantile(SME, probs=c(0.25, 0.75), na.rm=T)[1],
            SME_U = quantile(SME, probs=c(0.25, 0.75), na.rm=T)[2],
            # SME_L_50 = quantile(SME, probs=c(0.25, 0.75), na.rm=T)[1],
            # SME_U_50 = quantile(SME, probs=c(0.25, 0.75), na.rm=T)[2],
            # SME_L_95 = quantile(SME, probs=c(0.025, 0.975), na.rm=T)[1],
            # SME_U_95 = quantile(SME, probs=c(0.025, 0.975), na.rm=T)[2],
            SME = mean(SME, na.rm = T), 
            CV = unique(CV), 
            SRMSE = unique(SRMSE),
            i = unique(i))  %>%
  ungroup() %>%
  mutate(SRMSE = ifelse(oSCR_model == "d." & Parameter == "d.beta", NA, SRMSE)) %>%
  mutate(design = factor(design, levels = c("2sigma", "pbar", "p2bar", "pcombo"))) %>%
  filter(Parameter %in% c("p0", "sig", "EN"))  %>%
  drop_na(SME) %>%
  select(i, everything()) %>%
  arrange(i) %>%
  as.data.frame()


add1 = expand.grid(i = 100,  geom ="regular", density = "uniform", 
                   design = c("2sigma", "pbar", "p2bar", "pcombo"), coverage = c("A", "B", "C"),  
                   oSCR_model = "ds", Parameter = c("p0", "sig", "d0", "d.beta", "EN"), 
                   SME_L = NA, SME_U = NA, SME = NA, CV = NA, SRMSE = NA)

add2 = expand.grid(i = 100,  geom ="irregular", density = c("uniform", "weak", "strong"),
                   design = c("2sigma", "pbar", "p2bar", "pcombo"), coverage = c("A", "B", "C"),  
                   oSCR_model = "ds", Parameter = c("p0", "sig", "d0", "d.beta", "EN"), 
                   SME_L = NA, SME_U = NA, SME = NA, CV = NA, SRMSE = NA)

add = rbind(add1, add2)

plot_df = evals %>%
  rbind(., add) %>%
  mutate(geom = recode(geom,
                       "regular" = "Regular",
                       "irregular" = "Irregular")) %>%
  mutate(density = recode(density,
                          "uniform" = "Uniform",
                          "weak" = "Weak",
                          "strong" = "Strong")) %>%
  mutate(design = recode(design,
                         "2sigma" = "2~sigma",
                         "pbar" = "Q[bar(p)]",
                         "p2bar" = "Q[bar(p)[m]]",
                         "pcombo" = "Q[bar(p)[c]]")) %>%
  mutate(design = factor(
    design, levels = c("2~sigma", "Q[bar(p)]", "Q[bar(p)[m]]", "Q[bar(p)[c]]"))) %>%
  filter(Parameter == "p0")  %>%
  mutate(coverage = recode(coverage, "A"="144", "B"="100", "C"="49")) %>%
  mutate(oSCR_model = as.factor(oSCR_model)) %>%
  mutate(plot_group =  interaction(design, oSCR_model)) %>%
  mutate(plot_group = factor(
    plot_group, levels = c("2~sigma.d.", "2~sigma.ds", 
                           "Q[bar(p)].d.", "Q[bar(p)].ds",
                           "Q[bar(p)[m]].d.", "Q[bar(p)[m]].ds",
                           "Q[bar(p)[c]].d.", "Q[bar(p)[c]].ds")))

"_____________________Figures_____________________"


"____BIAS____"

ggplot(data = plot_df, 
       aes(x = coverage, y = SME, group = plot_group, 
           color = plot_group, fill = plot_group, shape = plot_group)) +
  
  # Add 0 line & +/- 5% error lines
  geom_hline(yintercept = c(-5,5), color = "gray80", 
             size = 0.25) +
  geom_hline(yintercept = 0, color = "gray80", size = 0.5) +
  
  # Add error bars
  geom_errorbar(aes(ymin=SME_L, ymax=SME_U),
                position = position_dodge(0.9), #col = "gray70",
                size = 0.25, width = 0, na.rm = T) +
  
  # Add data points
  geom_point(position = position_dodge(0.9), 
             size = 0.8, stroke = 0.65, na.rm = T) +
  
  # # Adjust shapes
  scale_shape_manual(values=c(21, 21, 24, 24, 22, 22, 23, 23)) +
  #scale_shape_manual(values=rep(21, 8)) +
  
  # Adjust colors
  scale_color_manual(values = c("gray30", "gray30", ibm[3], ibm[3], ibm[1], ibm[1], ibm[2], ibm[2]))  +
  scale_fill_manual(values = c("white", "gray30", "white", ibm[3], "white", ibm[1], "white", ibm[2])) +
  
  # Break into facets
  facet_grid(density~geom, labeller = label_parsed,
             scales = "free_x", space = "free_x") +
  
  # Adjust labels
  xlab("Effort") + ylab("%RB") +
  
  # Adjust theme
  theme_bw() +
  theme(panel.grid = element_blank(),
        title = element_text(size=9),
        strip.text = element_text(size=9),
        axis.text.y = element_text(size=7),
        axis.text.x = element_text(size=9),
        legend.position = "none")


# SAVE PLOT
ggsave(plot = last_plot(), device = "jpg",
       dpi = 600, width = 5,  height = 4.5, units = "in",
       filename = "Figure3.jpg", path = getwd())


"____PRECISION____"

ggplot(data = plot_df, 
       aes(x = coverage, y = CV, group = plot_group, 
           color = plot_group, fill = plot_group, shape = plot_group)) +
  
  # Add data points
  geom_point(position = position_dodge(0.9), 
             size = 0.8, stroke = 0.65, na.rm = T) +
  
  # # Adjust shapes
  scale_shape_manual(values=c(21, 21, 24, 24, 22, 22, 23, 23)) +
  #scale_shape_manual(values=rep(21, 8)) +
  
  # Adjust colors
  scale_color_manual(values = c("gray30", "gray30", ibm[3], ibm[3], ibm[1], ibm[1], ibm[2], ibm[2]))  +
  scale_fill_manual(values = c("white", "gray30", "white", ibm[3], "white", ibm[1], "white", ibm[2])) +
  
  # Break into facets
  facet_grid(density~geom, labeller = label_parsed,
             scales = "free_x", space = "free_x") +
  
  # Adjust labels
  xlab("Effort") + ylab("CV") +
  
  # Adjust theme
  theme_bw() +
  theme(panel.grid = element_blank(),
        title = element_text(size=9),
        strip.text = element_text(size=9),
        axis.text.y = element_text(size=7),
        axis.text.x = element_text(size=9),
        legend.position = "none")


# SAVE PLOT
ggsave(plot = last_plot(), device = "jpg",
       dpi = 600, width = 5,  height = 4.5, units = "in",
       filename = "CVplot.jpg", path = getwd())
```
